# Lipoic acid biosynthesis is essential for *Plasmodium falciparum* transmission and influences redox response and carbon metabolism of parasite asexual blood stages

**DOI:** 10.1101/2020.05.17.099630

**Authors:** Marco Biddau, T.R. Santha Kumar, Philipp Henrich, Larissa M. Laine, Gavin J. Blackburn, Achuthanunni Chokkathukalam, Tao B Li, Kim Lee Sim, Stephen L Hoffman, Michael P. Barrett, Graham H. Coombs, Geoffrey I. McFadden, David A. Fidock, Sylke Müller, Lilach Sheiner

## Abstract

Malaria is still one of the most important global infectious diseases. Emergence of drug resistance and a shortage of new efficient anti-malarials continue to hamper a malaria eradication agenda. Malaria parasites are highly sensitive to changes in redox environment. Understanding the mechanisms regulating parasite redox could contribute to the design of new drugs. Malaria parasites have a complex network of redox regulatory systems housed in their cytosol, in their mitochondrion and in their plastid (apicoplast). While the roles of enzymes of the thioredoxin and glutathione pathways in parasite survival have been explored, the antioxidant role of α-lipoic acid (LA) produced in the apicoplast has not been tested. We analysed the effects of LA depletion on mutant *Plasmodium falciparum* lacking the apicoplast lipoic acid protein ligase B (*lipB*). Our results showed a change in expression of redox regulators in the apicoplast and the cytosol. We further detected a change in parasite central carbon metabolism, with LA depletion influencing glycolysis and tricarboxylic acid cycle activity. Importantly, abrogation of LipB impacted *P. falciparum* mosquito development, preventing oocyst maturation and production of infectious sporozoite stages, thus flagging LA biosynthesis as a potential target for the development of new transmission drugs.

## INTRODUCTION

Malaria remains a tremendous threat to human health with 200 million infections resulting in 405,000 deaths in 2018 (World Health Organization, 2019), so it is imperative that we identify new antimalarial targets. One potential target are the parasite redox regulation systems. *Plasmodium falciparum* is constitutively exposed in all stages of its complex life cycle to molecules that challenge its redox balance. Finding ways to disrupt this delicate balance hold promise for drug development. Indeed, *in vitro* experiments in which parasites were exposed to exogenous H_2_O_2_-generating systems proved lethal for intra-erythrocytic stages (Dockrell and Playfair, 1984). Likewise, mature gametocytes are sensitive to the oxidative stress generated by exposure to redox-cyclers *in vitro* (Siciliano et al., 2017). Finally, an animal diet that generates an environment rich in reactive oxygen species (ROS) in hepatocytes hosting parasite cells resulted in reduced *Plasmodium* infection *in vivo* (Zuzarte-Luís et al., 2017). Despite this importance, and the fact that redox regulation is a fundamental aspect of cellular functions, our understanding of the parasite redox regulatory networks remains limited (Kehr et al., 2010; Müller, 2015).

The apicoplast, a non-photosynthetic plastid acquired via secondary endosymbiosis of a red algal cell, is an active metabolic hub in apicomplexan parasites including *Plasmodium* spp., (Biddau and Sheiner, 2019; Frohnecke et al., 2015; Kimata-Ariga et al., 2018; Mohring et al., 2014; Sheiner et al., 2013). The apicoplast has key roles in redox balance and hosts components of the thioredoxin and the glutathione systems, which represent the two best characterized cellular antioxidant systems. Apicoplast based redox regulators include the peroxiredoxin antioxidant protein (AOP), the dually-targeted (cytosol and apicoplast) enzymes glutathione reductase (GR), and the glutathione peroxidase-like thioredoxin peroxidase (TPx_Gl_) (Kehr et al., 2010; Laine et al., 2015). Likewise, two glyoxalase system proteins are apicoplast targeted: glyoxalase-1-like protein (GILP), and glyoxalase 2 (tGloII) (Kehr et al., 2010; Urscher et al., 2010). Glyoxalase 2 is proposed to play a role in detoxification of incomplete triosephosphate-isomerase reaction products, but is apparently dispensable during intra-erythrocytic development (Wezena et al., 2017). Two additional apicoplast thioredoxin-like proteins (ATrx1 and ATrx2) are found in the peripheral compartments (Sheiner et al., 2011). ATrx1 and ATrx2 were recently characterised in *T. gondii* and shown to play an essential role in the control of protein sorting and folding in response to organelle redox status (Biddau et al., 2018). The *Plasmodium* orthologue of ATrx2 (*Pf*ATrx2; PF3D7_0529100) is likely essential too (Bushell et al., 2017; Zhang et al., 2018). Many other potential redox-active proteins are predicted to be localised in the apicoplast (Boucher et al., 2018), but their roles are uncharacterized.

An additional molecule proposed to take part in apicoplast redox regulation is α-lipoic acid (LA) (Frohnecke et al., 2015; Günther et al., 2007; Laine et al., 2015). Due to its reducing properties, LA is known as the ‘universal antioxidant’ (Gorąca et al., 2011; Kagan et al., 1992; Moura et al., 2015; Perham, 2000; Tibullo et al., 2017). The proposed antioxidant role of LA in the apicoplast is based on a link between redox regulation and apicoplast pyruvate metabolism via the pyruvate dehydrogenase enzyme complex (PDC). The three enzymes in the PDC complex are pyruvate dehydrogenase (E1), dihydrolipoyl transacetylase (E2) and apicoplast dihydrolipoyl dehydrogenase (aE3). Through a series of reactions, PDC transfers an acetyl group from pyruvate to coenzyme A (CoA) to generate acetyl-CoA for the fatty acid biosynthesis pathway (Foth et al., 2005; Mooney et al., 2002) (Fig. 1). This activity depends on LA bound to the E2 lipoyl domain, which is reduced to dihydrolipoic acid (DHLA) during the process. The final reaction of *Plasmodium* PDC is catalysed by aE3, which re-oxidises DHLA back to LA to allow another cycle of PDC activity (Fig. 1). The activity of aE3 is coupled to the reduction of NAD^+^ to NADH+H^+^, which in turn takes part in apicoplast redox regulation (Laine, 2014; McMillan et al., 2005). The DHLA/LA redox couple has a redox potential of −0.32 V, which is lower than the glutathione/glutathione disulphide (GSH/GSSG) couple potential of −0.24 V, thus making glutathione a potential substrate for DHLA (Packer et al., 1995).

**Figure 1.**
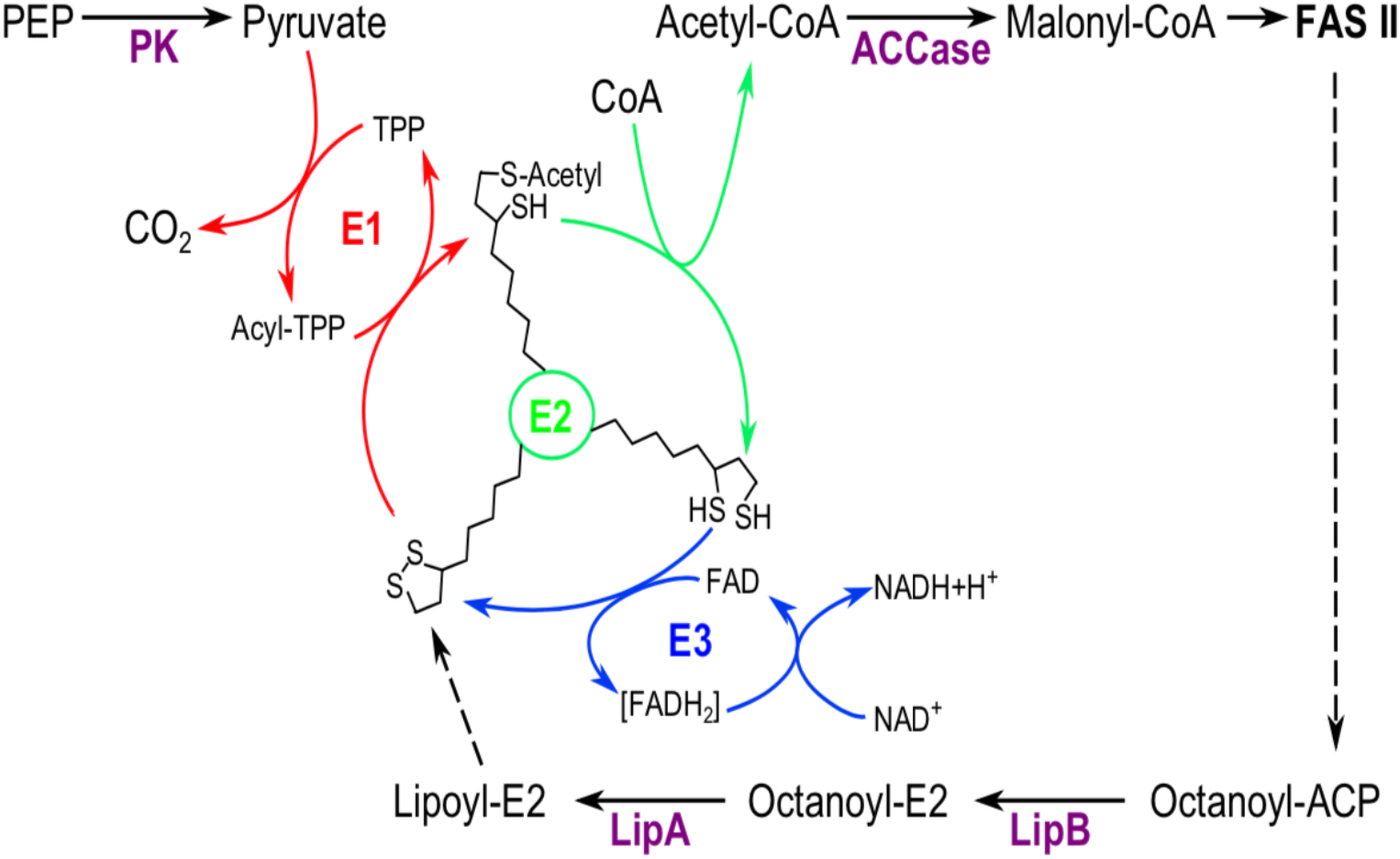
Schematic representation of PDC and LA biosynthesis components and function in *P. falciparum* apicoplast. Each of the PDC enzyme reactions is shown in different colour (E1, red; E2, green; E3, blue). The three states of E2-conjugated LA are depicted onto the schematic E2. Abbreviations: ACCase: Acetyl-CoA carboxylase; CoA: Coenzyme A; FAD: Flavin adenine dinucleotide; FAS II: Fatty acid biosynthesis type II; LipA: lipoyl synthase; LipB: Octanoyl-ACP:protein *N*-octanoyltransferase; NAD: Nicotinamide adenine dinucleotide; PK: Pyruvate kinase; TPP: Thiamine pyrophosphate.

The apicoplast retains its exclusive LA biosynthesis pathway, catalysed by the enzymes octanoyl-ACP:protein *N*-octanoyltransferase (LipB) and lipoyl synthase (LipA)(Fig. 1), which operates independently from the mitochondrial LA salvage (Crawford et al., 2006; Günther et al., 2009). To test the putative role of LA in redox regulation, and its link to parasite metabolism, we examined the changes in the expression of redox regulation enzymes and the metabolic changes occurring in a *P. falciparum* mutant (named 3D7^Δ*PflipB*^) where the *lipB* gene is disrupted (Günther et al., 2007). As a comparison, we also analysed an aE3 deletion mutant that has only a mild effect on redox balance in the parasite (Laine et al., 2015), potentially because aE3 function might be compensated by alternative apicoplast enzymatic systems coupled to NAD(P)^+^ reduction such as the NADP^+^-specific glutamate dehydrogenase (Zocher et al., 2012). Our results support a link between LA availability and redox regulation. Additionally, LipB depletion led to changes in central carbon metabolism corroborating a link between apicoplast redox regulation and cytosolic and mitochondrial metabolic pathways.

Importantly, LipB depletion hampers the ability of the parasites to develop in the mosquito, which is in line with the crucial functions of redox regulation and fatty acid biosynthesis in the insect stage of the parasite life cycle (Cobbold et al., 2013; Pastrana-Mena et al., 2010; Siciliano et al., 2017; van Schaijk et al., 2014).

## RESULTS

### Deletion of *lipB* modulates apicoplast and cytosol antioxidant levels

Pronounced LA deficiency was earlier detected in the trophozoite stage in LipB KO line 3D7^Δ*PflipB*^ parasites (Günther et al., 2007). To investigate whether this deficiency affected the apicoplast antioxidant composition, we monitored the relative transcription levels of genes encoding apicoplast antioxidants at 26-, 30- and 34-hours post-invasion (hpi) by qPCR for both 3D7^Δ*PflipB*^ and the parental (3D7^WT^) parasites. We tested the apicoplast redox-active proteins TPx_Gl_, ATrx2 and AOP (Fig. 2), which take part in the thioredoxin redox system, and the apicoplast glyoxalase system proteins GILP and tGlo (Fig. S1). Among these, TPx_Gl_ had the most dramatic change in expression levels. Relative transcription levels of TPx_Gl_ displayed three to four-fold increases at 30 hpi and 34 hpi when compared to 3D7^WT^. Similarly, ATrx2 relative expression showed a four-fold increase compared to the 3D7^WT^ parasites at 26 hpi followed by a two-fold decrease at 34 hpi, when AOP also showed 2-fold decrease (Fig. 2). In contrast, the apicoplast glyoxalase system enzymes presented no significant differences in relative expression (Fig. S1).

**Figure 2.**
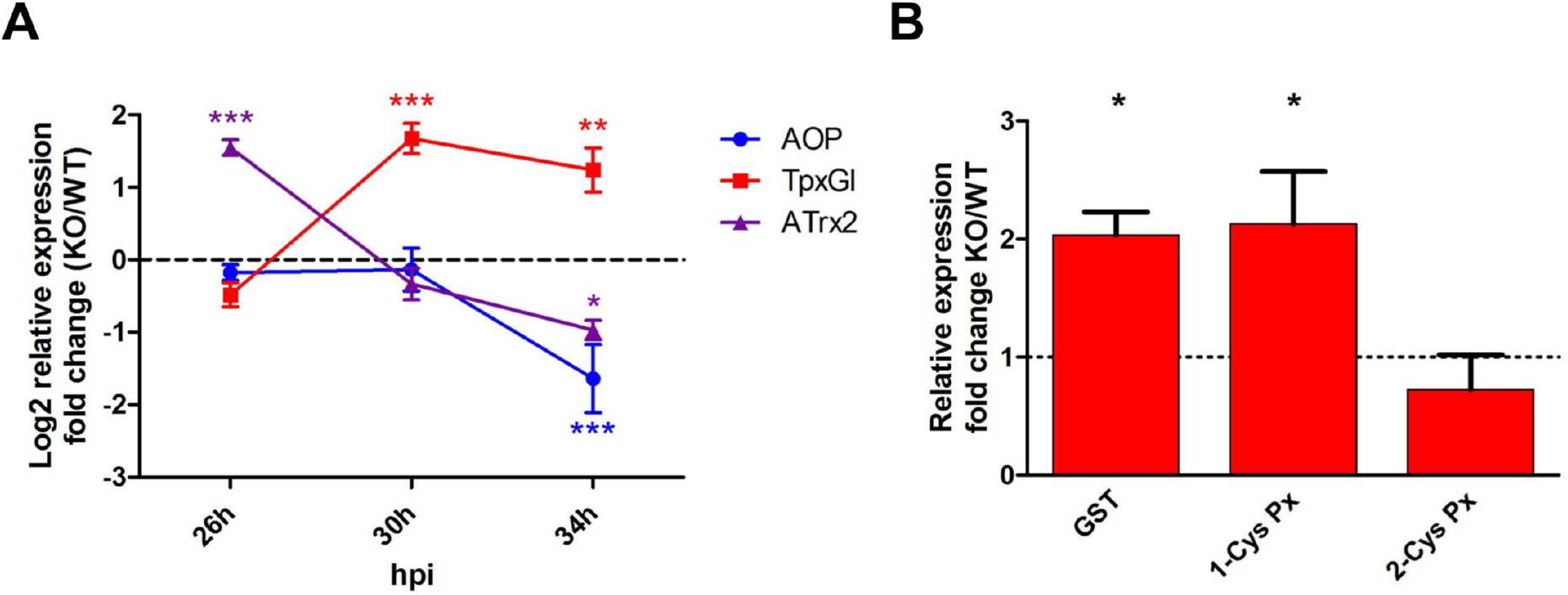
Analysis of apicoplast antioxidant relative expression levels and cytosolic antioxidant relative protein levels. (**A**) Relative expression levels for apicoplast antioxidant protein (AOP), glutathione peroxidase-like thioredoxin peroxidase (TPx_Gl_), and apicoplast thioredoxin-like protein 2 (ATrx2), in the LipB mutant compared to wild type. Parasites were highly synchronised following the sorbitol and MACS protocol (see Materials and Methods) and harvested at 26, 30 and 34 hours post-invasion (hpi). Differences are expressed as Log_2_ of the 3D7^Δ*PflipB*^/3D7^WT^ ratio of the mean signals from three experiments performed in biological triplicates (n=3). Error bars show SD. Variances were analysed using the 2-way ANOVA test coupled with the Bonferroni test using GraphPad Prism 5. Asterisks and graph lines are colour coded as shown in the legend; *: P<0.05; **: P<0.01; ***: P<0.001. (**B**) Relative protein levels for the cytosolic antioxidant proteins glutathione S-transferase (GST), 1-Cys peroxiredoxin (1-CysPx) and 2-Cys peroxiredoxin (2-CysPx). Three independent experiments were performed in biological triplicates and the means (n=3) of actin-normalised fluorescent signals for each protein were calculated using quantitative fluorescent western blotting. The bars represent the 3D7^Δ*PflipB*^/3D7^WT^ average ratio ± SD. The variance was analysed with the Student t-test using GraphPad Prism 5; *: P<0.05.

In light of this apicoplast antioxidant response, we wanted to test whether the cytosolic antioxidant composition was also affected. Therefore, we used quantitative fluorescent western blotting to monitor the relative levels of glutathione S-transferase (GST; PF3D7_1419300), 1-Cys peroxiredoxin (1CysPx; PF3D7_0802200) and 2-Cys peroxiredoxin (2-CysPx; PF3D7_1438900). These proteins were measured in 3D7^Δ*PflipB*^ and 3D7^WT^ late trophozoites at 34 hpi. We observed a significant two-fold increase in protein levels for GST and 1-CysPx in 3D7^Δ*PflipB*^ mutants (Fig. 2B). Conversely, 2-CysPx levels appeared to remain unchanged between the two lines (Fig. 2B).

### Deletion of *lipB* affects parasite carbon metabolism

In eukaryotic cells, compartmental redox state and the availability of redox conducting and regulating molecules in different cellular compartments are intertwined with the activity of metabolic pathways. The redox conditions in a cellular compartment affect the function of its metabolic enzymes, while in return, the metabolic reactions in a compartment generate metabolites that impact its redox state. Thus, we proceeded to examine whether the observed redox changes in 3D7^Δ*PflipB*^ mutants coincide with changes in central carbon metabolism. We chose to make this analysis alongside a *P. falciparum* apicoplast dihydrolipoamide dehydrogenase (aE3) knock-out mutant (3D7^Δ*Pfae3*^) (Laine et al., 2015). Unlike 3D7^Δ*PflipB*^, the deletion of a PDC component in 3D7^Δ*Pfae3*^ does not result in disruption of LA biosynthesis nor of PDC activity and its effect on the expression of redox regulators is only mild (Laine et al., 2015), which makes 3D7^Δ*Pfae3*^ an ideal negative control.

The levels of D-glucose and L-lactate in spent medium were monitored using commercial enzymatic assays in two independent experiments. As glycolytic activity in *P. falciparum* typically peaks during intra-erythrocytic trophozoite development (Shivapurkar et al., 2018), we collected samples at 30, 34, 38 and 42 hpi to cover this developmental stage. Results showed that spent medium samples from 3D7^Δ*PflipB*^ contained significantly less D-glucose and more L-lactate than 3D7^WT^ at 42 hpi (Fig. 3). Conversely, 3D7^Δ*Pfae3*^ mutants did not present this trend and the concentrations for these metabolites in spent medium were comparable to the WT controls (Fig. S2).

**Figure 3.**
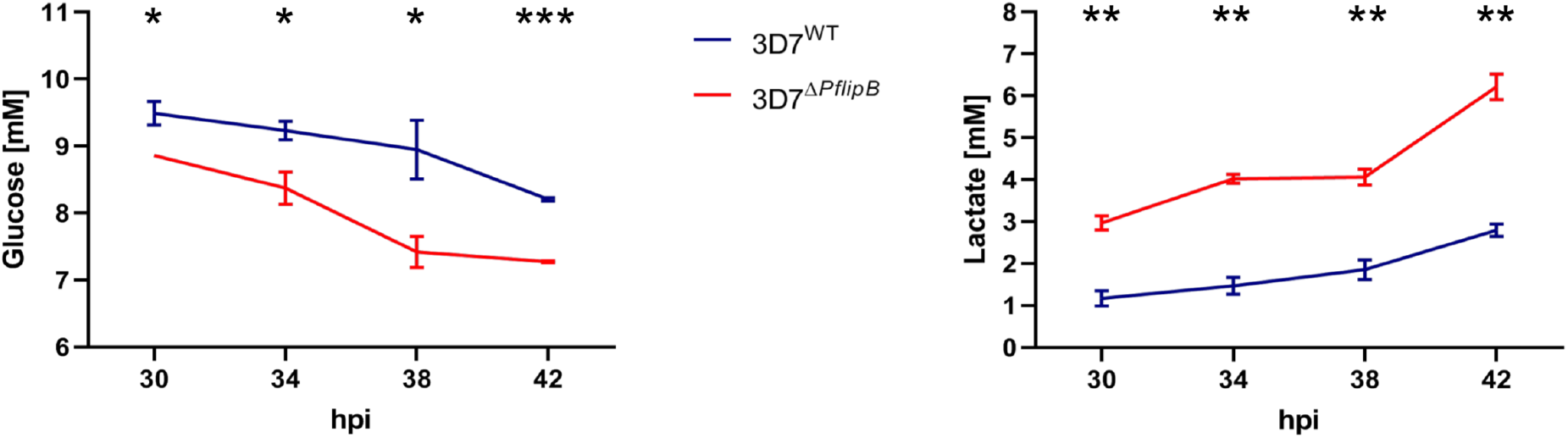
Analysis of D-glucose (left) and L-lactate (right) in spent medium samples from 3D7^Δ*PflipB*^ mutants and 3D7^WT^ parasites cultures. Spent medium samples were collected at 30, 34, 38 and 42 hpi and analysed using a commercial enzymatic assay for D-glucose and L-lactate. Triplicate cultures at 2% parasitemia were used for each parasite line for this experiment. The mean signals from 2 independent experiments (n=2) are reported with error bars representing SD. The variance between the lines at each time point was analysed with the Student t-test using GraphPad Prism 5; *: P<0.05; **: P<0.01; ***: P<0.001.

We further hypothesised that the up-regulation of glycolysis and the change to antioxidant expression described above might affect downstream metabolism and, especially the tricarboxylic acid (TCA) cycle. Therefore, we proceeded to set up two steady-state targeted metabolomics experiments in biological triplicates, using the isotope-labelled nutrient ^13^C-U-D-glucose. 3D7^Δ*PflipB*^ and 3D7^WT^ parasites were synchronised and metabolically labelled for 28 hours. Parasites at the late trophozoite stage were then rapidly chilled, and the extracted metabolites analysed by liquid chromatography-mass spectrometry (LC-MS) to follow [^13^C] labelling. The analysis of the labelled fraction for these metabolites showed an immediate conversion of glucose into glycolytic intermediates (Fig. S3), in line with previous analyses (Storm et al., 2014). In agreement with the observation from the spent medium, 3D7^Δ*PflipB*^ mutants displayed a two-fold increase in the relative levels of the glycolytic metabolite pyruvate and of glycerol-3-phosphate, which derive from a glycolytic intermediate (Fig. 4A, B). A similar trend was also displayed by the metabolites 2-oxoglutarate and succinate associated with the TCA cycle, as well as the amino acid aspartate, whereas alanine displayed the reverse tendency (Fig. 4A, B). Conversely, the analysis of 3D7^Δ*Pfae3*^ mutants showed no significant differences in the relative abundances for these metabolites (Fig. 4B).

**Figure 4.**
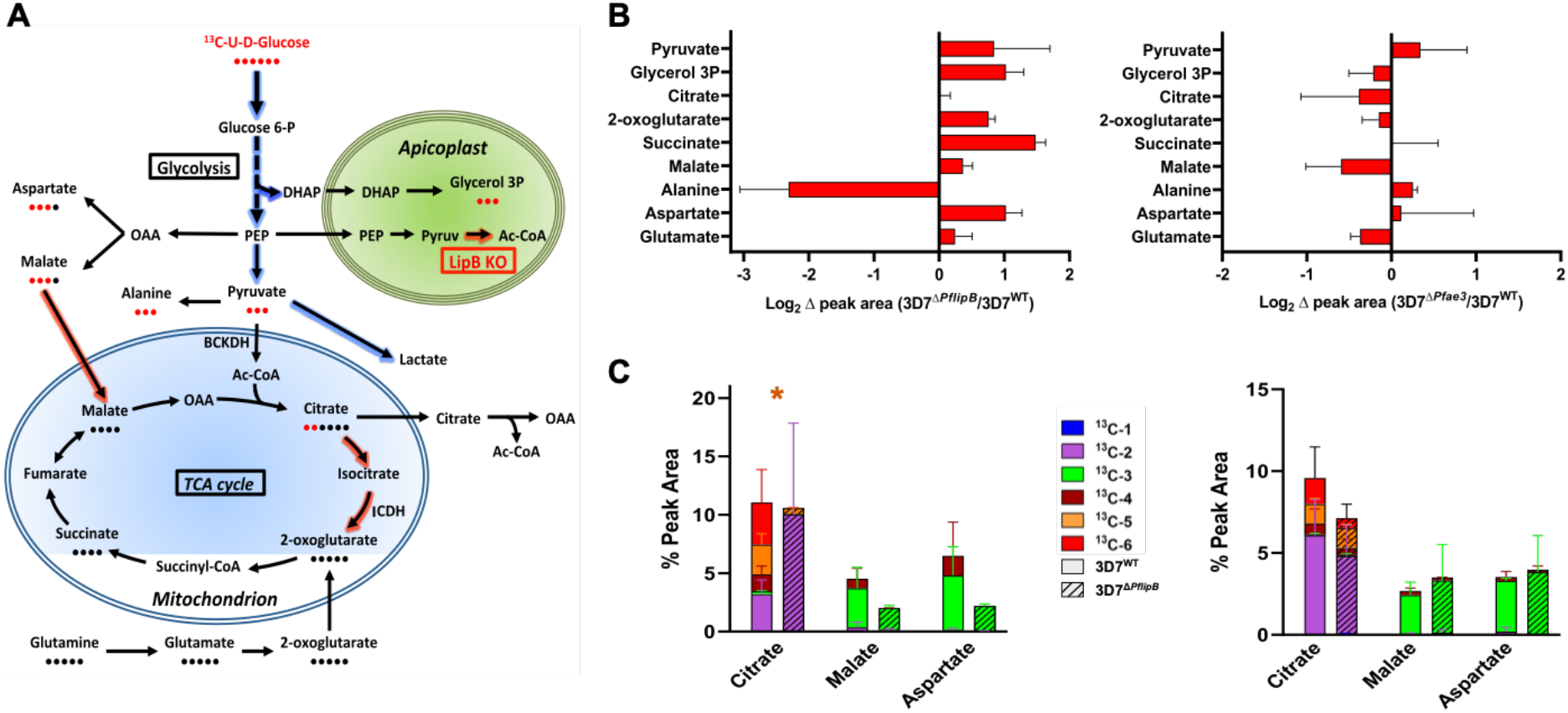
Metabolomic analyses of 3D7^Δ*PflipB*^ mutants and 3D7^WT^ parasites using ^13^C-U-D-glucose labelling. Results from two independent targeted metabolomics experiments in biological triplicates comparing 3D7^Δ*PflipB*^ mutants and 3D7^WT^ after incubation in culture medium containing 100% ^13^C-U-D-glucose for 28 hours. (**A**) Schematics of the *P. falciparum* central carbon metabolism pathways analysed here, highlighting the metabolic adaptations in 3D7^Δ*PflipB*^ mutant compared to wild-type parasites. Arrows shaded in red and blue respectively correspond to decreased and increased in flux for each specific reaction. Red and black dots under metabolite names respectively depict the number of labelled and unlabelled carbons, based on the most abundant labelled form of the metabolite. Abbreviations: Ac-CoA: Acetyl-CoA; BCKDH: branched-chain ketoacid dehydrogenase; DHAP: Dihydroxyacetone phosphate; OAA: oxaloacetate; PEP: Phospho*enol*pyruvate. (**B**) Relative intracellular levels for each metabolite obtained by the sum of all the peak areas for each isotopologue of a specific metabolite. The differences in abundance for each metabolite are expressed as Log_2_ of the 3D7^Δ*PflipB*^/3D7^WT^ (left graph) or Log_2_(3D7^Δ*Pfae3*^/3D7^WT^) (right graph) mean ratio ± SD (n=2). (**C**) Bar graphs summarising the percentage of isotopic incorporation in each identified metabolite calculated from the chromatographic peak areas. The bars are divided to represent the mean contribution (n=2) of the different isotopologues to each metabolite labelled fraction and error bars are SD. Empty bars represent metabolites identified in 3D7^WT^ parasites, while dashed bars correspond to metabolites from 3D7^Δ*PflipB*^ mutants (left graph) or 3D7^Δ*Pfae3*^ mutants (right graph). The variance of citrate M+5 fraction between mutants and 3D7^WT^ is depicted by an asterisk coloured with the corresponding legend colour and was analysed with the Student t-test using GraphPad Prism 5; *: P<0.05.

Only a small fraction of [^13^C] labelled triose phosphates was fed into the TCA cycle intermediates in all three parasite lines (Fig. S3), as has been well established in the literature (Ke et al., 2015; MacRae et al., 2013; Storm et al., 2014). Interestingly, whereas the 3D7^WT^ and 3D7^Δ*Pfae3*^ controls had M+4, M+5 and M+6 citrate labelling, indicative of a complete TCA cycle activity (Fig. 4C), this was not the case for 3D7^Δ*PflipB*^. Rather, in 3D7^Δ*PflipB*^ the signals for M+4 and M+6 citrate were below the detection level, while the M+5 fraction was significantly decreased (Fig. 4C). In both experiments an increment of M+2 citrate was detected for 3D7^Δ*PflipB*^ mutants compared to 3D7^WT^, however, with high variability between the two experiments (Fig. 4C). This labelling pattern, suggested a potential altered flux of glycolytic carbons into the TCA cycle. We further tested this possibility by quantitative western blot analysis of the enzymes operating the first steps of the TCA cycle. Interestingly, the 3D7^Δ*PflipB*^ mutant up-regulate the branched-chain α-keto acid dehydrogenase (BCKDH) component E2, the first enzyme converting pyruvate to acetyl co-A (Oppenheim et al., 2014), while its apicoplast parallel PDC E2 displayed no differences in abundance (Fig. 5A). In contrast, isocitrate dehydrogenase (ICDH), which operates three steps downstream in the TCA cycle, displayed no differences in abundance when compared with 3D7^WT^ (Fig. 5A). Collectively these results point to an altered flux of metabolites through the TCA cycle such that glycolytic derived carbon provide less input while glutamate becomes a major TCA cycle carbon input.

**Figure 5.**
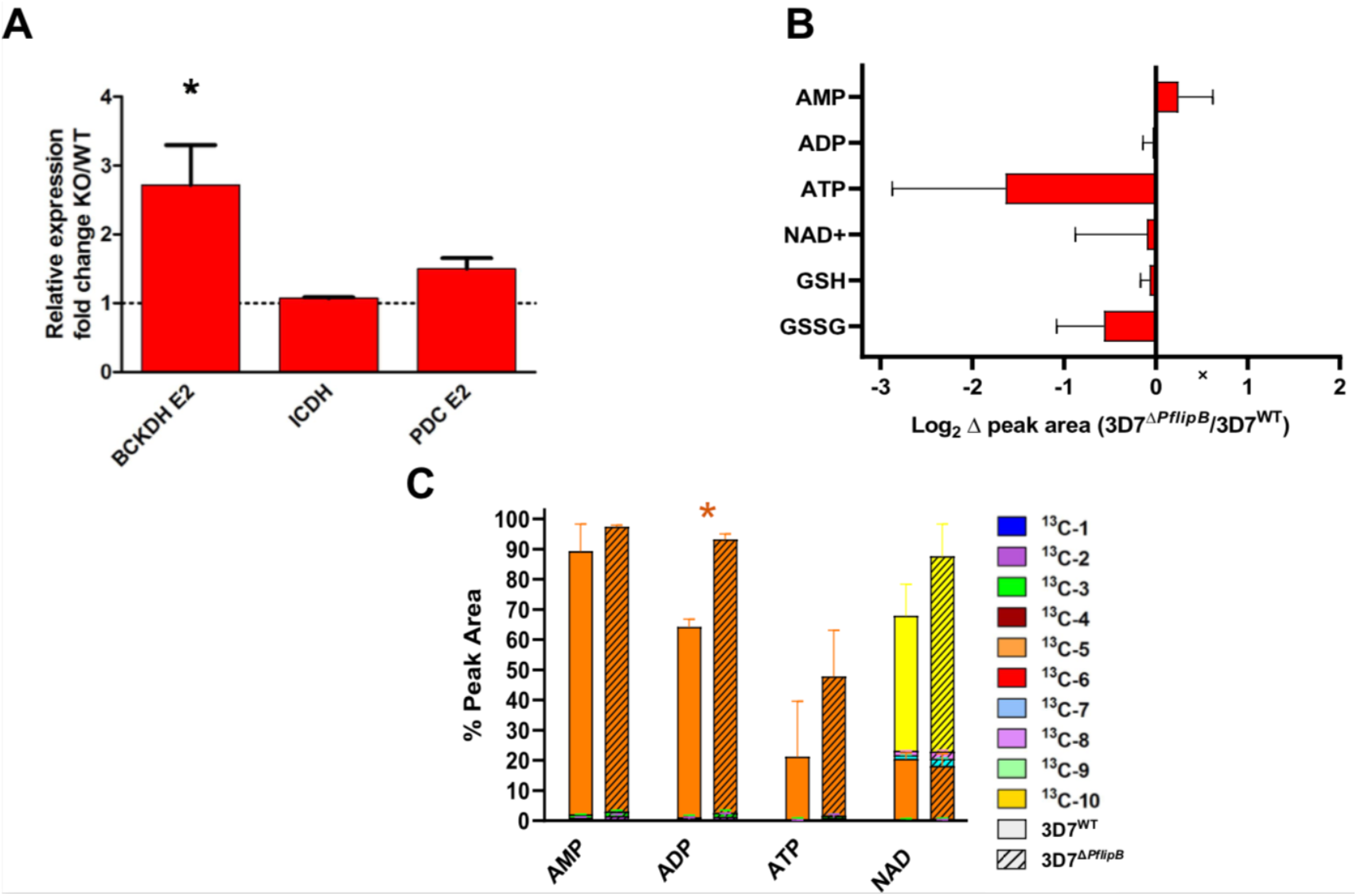
Analysis of the expression of metabolic enzymes and the levels of cofactors involved in glycolysis and in the TCA cycle. (**A**) Relative protein levels for the mitochondrial enzymes branched-chain ketoacid dehydrogenase (BCKDH E2), isocitrate dehydrogenase (ICDH) and of the apicoplast enzyme dihydrolipoamide transacetylase (PDC E2). Data from three experiments performed in biological triplicates (n=3) are shown as 3D7^Δ*PflipB*^/3D7^WT^ ratio of the actin-normalised mean fluorescent signals for each protein. Error bars represent SD. The variance was analysed with the Student t-test using GraphPad Prism 5; *: P<0.05. (**B**) Relative intracellular levels of metabolic cofactors are represented as Log_2_ of the 3D7^Δ*PflipB*^/3D7^WT^ ratio of the means from two experiments in biological triplicates. The relative mean levels are the sum of all peak areas relative to each isotopologue of each metabolite (n=2). Error bars represent SD. (**C**) Bar graph summarising the percentage of isotope incorporation in the identified cofactors AMP, ADP, ATP and NAD^+^. Data correspond to two experiments performed in biological triplicates (n=2). The bars are divided to the mean contribution of each isotopologue to the total labelled fraction and displays error bars correspond to SD. Empty bars correspond to metabolites identified in 3D7^WT^ parasites, while dashed bars refer to metabolites from 3D7^Δ*PflipB*^ mutants. The variance of each isotopologue fraction between 3D7^Δ*PflipB*^ and 3D7^WT^ (n=2) is resented by the asterisk symbol (coloured according to legend) and was analysed with the Student t-test using GraphPad Prism 5; *: P<0.05.

Lastly, we tested cofactors involved in carbon and energy metabolism, where we observed a decrease in relative intracellular levels of ATP in 3D7^Δ*PflipB*^, while all other cofactors tested were unchanged (Fig. 5B). In addition, 3D7^Δ*PflipB*^ mutants had an increase in the ADP M+5 fraction (Fig. 5C). These results may suggest an increased ATP demand in the mutants. Likewise, an observed increase in incorporation of glucose-derived fully labelled ribose, contributing to the M+5 fraction of this metabolite, may point to an up-regulation of ADP generation through the addition of the ribose to salvaged hypoxantine.

In summary, the analysis of 3D7^Δ*PflipB*^ mutant metabolism reveals an effect on the activity of both glycolysis and the TCA cycle. The specificity of this phenotype compared to 3D7^Δ*Pfae3*^ metabolism supports the role of LA biosynthesis in cellular homeostasis.

### Can the *lipB* mutant complete mosquito development?

Despite the observed changes in redox regulation and metabolic fluxes, 3D7^Δ*PflipB*^ growth in RBC culture was only mildly affected, showing a slight increase in growth (Günther et al., 2007). Likewise, we found that at 30 hpi the 3D7^Δ*PflipB*^ mutant showed an increase in differentiation into schizonts with a 6-hour advance compared to 3D7^WT^ (Fig. S4A). This accelerated differentiation resulted in faster completion of the trophozoite stage and of the whole asexual cycle but had no effect on continuous growth in culture and no differences in the average number of merozoites (Fig. S4B). We thus investigated the development *of lipB*-deleted parasites in mosquitoes. For this analysis, a second *lipB* deletion line was generated by double-crossover gene deletion (Fig. S5) in the NF54 background (NF54^Δ*PflipB*^), which unlike 3D7 parasites, is able to infect mosquitoes. Parasites were maintained in media containing 10% human serum, with the aim of them retaining their capacity to form mature gametocytes. The deletion of *lipB* did not appear to overtly influence sexual commitment in the parasites and mature gametocytes developed as in NF54^WT^ (Fig. 6, Table 1).

**Figure 6.**
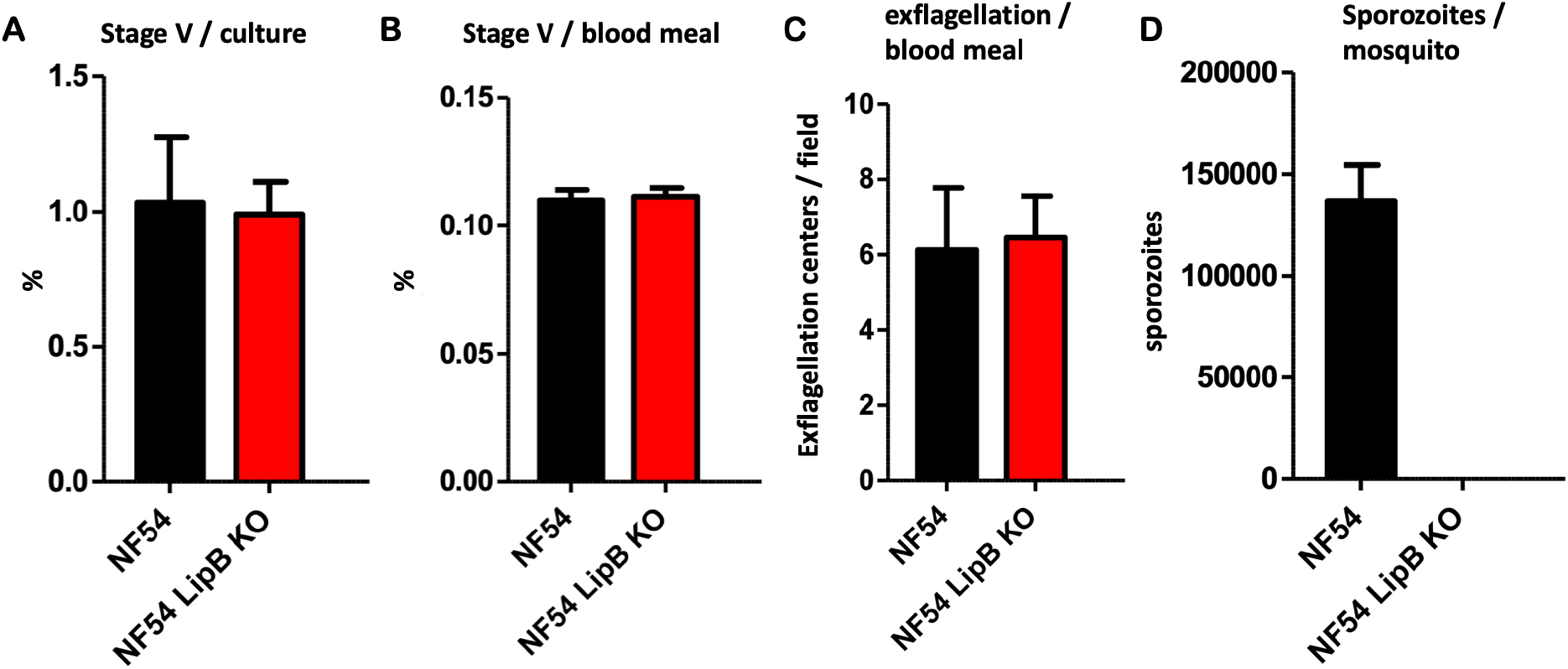
Bar graphs representing the data from Table 1.

**Table 1:** Summary of mosquito infection attempts with NF54^*ΔPflipB*^.

| Stage V gametocytemia in cultures |  | Final concentration of stage V gametocytemia in the infectious blood meals |  | Wet mount exflagellation of infectious blood meal per field (x40 phase contrast) |  | Sporozoites/ mosquito |  |
| --- | --- | --- | --- | --- | --- | --- | --- |
| NF54 | NF54 LipB KO | NF54 | NF54 LipB KO | NF54 | NF54 LipB KO | NF54 | NF54 LipB KO* |
| 0.40% | 1.62% | 0.11% | 0.11% | 5.2 | 11.7 | 106,543 | 0 |
| 1.37% | 0.63% | 0.12% | 0.12% | 8.1 | 6.3 | 150,178 | 0 |
| 1.45% | 0.75% | 0.11% | 0.11% | 9.3 | 4.9 | 180,991 | 0 |
| 0.92% | 0.93% | 0.10% | 0.10% | 1.9 | 2.7 | 109,975 | 0 |
| -- | 0.87% |  | 0.10% |  | 9.4 |  | 0 |
| -- | 1.10% |  | 0.12% |  | 5 |  | 0 |
| -- | 1.03% |  | 0.12% |  | 5.6 |  | 0 |
| 1.04% ± 0.24% | 0.99% ± 0.12% | 0.11% ± 0.004% | 0.11% ± 0.003% | 6.1 ± 1.7 | 6.5 ± 1.2 | 136,922 ± 17,717 | 0 ± 0 |

Each row represents a separate experiment, with parental NF54 included in the first four. Despite equivalent numbers of mature gametocytes and rates of exflagellation of male gametes in comparison with parental NF54 parasites, the LipB KO parasites failed to produce any salivary gland sporozoites, as determined from dissection and counting of 20 infected *Anopheles* mosquitoes per parasite line per experiment. The bottom row shows composite mean±SEM data.

This allowed us to proceed to evaluate parasite development in the mosquito. Seven mosquito infection experiments were performed with NF54^Δ*PflipB*^ gametocytes (Table 1). While midgut oocysts were detectable in all experiments, no sporozoites were detected in any of the infected mosquitoes in any of the seven experiments, while an average of 136,922 ± 17,717 sporozoites was detected in four experiments performed with the parental NF54 (Table 1, Fig. 6). These data indicated a major defect in development in the mosquito in the NF54^Δ*PflipB*^ parasites. To explore this further, we examined the morphology of the midgut oocysts (Fig. 7). NF54^Δ*PflipB*^ parasites were unable to form midgut oocyst sporozoites, suggesting an attenuated sexual development for this line that could not complete its transmission cycle in the *Anopheles* vector.

**Figure 7.**
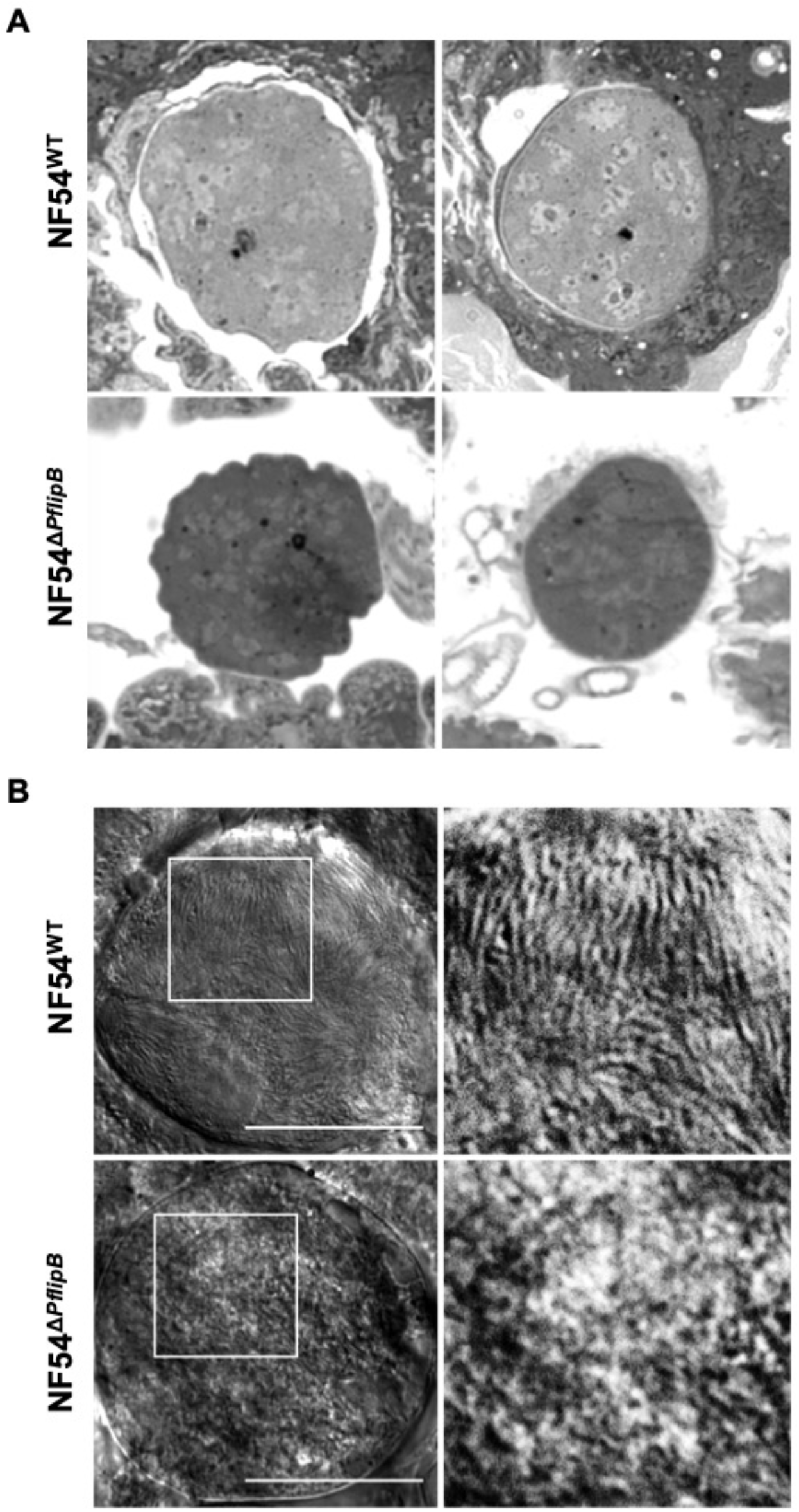
Oocyst morphology on day 7 and 13 post-feeding of gametocytes to *Anopheles* mosquitoes shows a defect in sporozoite development for LipB knockout parasites. (**A**) Two representative light microscopy images of NF54^Δ*lipB*^ and of NF54^WT^ oocytes on day 7 post feeding. (**B**) A representative DIC microscopy image of NF54^Δ*lipB*^ and of NF54^WT^ oocytes on day 13. The insets are shown in white squares. Scale bar 25 µm. The NF54^Δ*PflipB*^ oocysts show malformation in both methods and time points.

## DISCUSSION

The cellular redox balance of *Plasmodium* parasites is constantly under threat of oxidative stress generated by the metabolic functions of the parasite and by the metabolic activities and defence mechanisms of the host (Becker et al., 2004; Müller, 2015; Nepveu and Turrini, 2013; Patzewitz et al., 2013). Apicoplast-specific redox balance is an integral part of the overall cellular redox steady-state (Biddau et al., 2018; Kehr et al., 2010; Mohring et al., 2017). While fragmented information is available about the different apicoplast redox control pathways, their importance is evident in the series of specific antioxidant systems it hosts (Kehr et al., 2010) and in the redox regulators controlling its biogenesis (Biddau et al., 2018). Likewise, apicoplast-hosted pathways are coupled to redox reactions, including the biosynthesis of isoprenoid precursors, which is coupled to the reduction of NADP^+^ to NADPH plus H^+^ (Seeber et al., 2005; Seeber and Soldati-Favre, 2010), and the activity of PDC as discussed here (Fig. 1). LA is proposed to contribute to redox regulation in other systems (Tibullo et al., 2017), and here we provide evidence in support of this role within the apicoplast of *Plasmodium*.

### Evidence supporting the proposed role of LA as an apicoplast redox regulator

LA is a powerful antioxidant with low redox potential (Bilska and Włodek, 2005; Packer et al., 1995), which prompted us to test how interfering with LA biosynthesis affected apicoplast redox regulation. Our results revealed that the LipB deletion mutant (3D7^Δ*PflipB*^) shows transcriptional changes of the apicoplast redox enzymes peroxidase-like enzyme TPx_Gl_, thioredoxin ATrx2, and the peroxiredoxin AOP (Fig. 2A). We propose that these changes promote apicoplast redox homeostasis in response to the oxidative stress caused by the depletion of the LA antioxidant function. In support of this hypothesis, up-regulation of TPx_Gl_ also occurs in response to other oxidative stresses in *P. falciparum* (Akide-Ndunge et al., 2009). Likewise, we recently reported that the ortholog of *Pf*ATrx2 in the related parasite *Toxoplasma gondii* (*Tg*ATrx2) controls apicoplast gene expression, likely via a redox state-controlled interaction with proteins in transit to the apicoplast lumen (Biddau et al., 2018). If *Pf*ATrx2 performs a similar function, then the changes in its expression in 3D7^Δ*PflipB*^ may serve to control protein transit to the apicoplast lumen in response to organelle redox imbalance. The different roles of TPx_GI_ and ATrx2 may account for the different pattern in their transcriptional changes.

Our observations provide evidence linking the depletion of apicoplast LA to changes in the apicoplast antioxidant response (Fig. 2A). We thus suggest that the apicoplast *Pf*PDC E2 enzyme may operate as an apicoplast antioxidant through its prosthetic LA. Examples of DHLA acting as an electron donor to both GSH and thioredoxin systems have been previously described in other organisms (Packer et al., 1995). DHLA bound to PDC-E2 and to α-ketoglutarate dehydrogenase (KGDH) acts as an electron donor to glutaredoxins in an *E. coli* mutant with both the thioredoxin and the GSH systems were disrupted (Feeney et al., 2011). Likewise, *Mycobacterium tuberculosis* KGDH E2 uses DHLA to transfer electrons received from E3 to peroxiredoxins and contributes to antioxidant defence (Bryk et al., 2002). Additionally, KADH E2-mediated reduction of thioredoxins was observed in mammals (Bunik and Follmann, 1993). A similar redox regulatory role of the apicoplast PDC E2 may explain why it is expressed during the intra-erythrocytic stages (Foth et al., 2005; McMillan et al., 2005), despite the dispensability of fatty acid biosynthesis in this stage (Vaughan et al., 2009). Our attempts to delete PDC E2 in *P. falciparum* using gene replacement were unsuccessful (data not shown), raising the possibility that its role during intra-erythrocytic development is essential. In agreement with this observation, *Pfpdce2* was proposed to be essential during intra-erythrocytic development in a recent whole-genome random mutagenesis screen (Zhang et al., 2018). This hypothetical role for LA in redox regulation raises questions about the sources of electrons for LA/DHLA recycling. This might not be attributed exclusively to the flux of glycolytic pyruvate to PDC as this was suggested to contribute to a build-up of acetyl-CoA in the organelle (Lim and McFadden, 2010). A possible alternative candidate for LA/DHLA recycling might be the apicoplast-targeted GSH reductase (GR) (Müller, 2015). The GR-mediated reduction of LA was demonstrated in rats, and, in particular, the use of NADPH +H^+^ as an electron donor was described in mitochondrial fractions (Haramaki et al., 1997; Pick et al., 1995).

### Changes in the expression of cytosolic redox regulators and in the asexual cycle progression upon *lipB* deletion suggest putative apicoplast-to-cytosol signalling

Our data identified changes in cytosolic antioxidant expression in 3D7^Δ*PflipB*^ parasites. Both enzymes for which we observed upregulation, GST and 1-CysPx (Fig. 2B), are highly abundant in the *P. falciparum* cytosol (Liebau et al., 2002), and both have demonstrated antioxidant activity in this compartment (Deponte and Becker, 2005; Harwaldt et al., 2002; Krnajski et al., 2001; Liebau et al., 2002). One potential explanation for this phenotype is plastid-to-cytosol redox signalling, as described in plant chloroplasts (reviewed in Dietz et al., 2016). In *Arabidopsis thaliana,* chloroplast-originated H_2_O_2_ signal induces upregulation of the expression of genes encoding different GSTs as well as enzymes involved in glycolysis and the pentose phosphate pathway (PPP) (Sewelam et al., 2014). Moreover, chloroplast-originated ROS signalling induce changes in the cell cycle progression (Bode et al., 2016). It is thus possible that the accelerated cell cycle progressions described before for 3D7^Δ*PflipB*^ (Günther et al., 2007) may be triggered in response to a redox signalling originating from the apicoplast. Furthermore, trophozoites appear to be the asexual stage most affected by redox imbalance in culture (Akide-Ndunge et al., 2009), which is in line with our observations of accelerated differentiation at that stage in the mutant (Fig. S4).

### Deletion of *lipB* affects energy metabolism in *P. falciparum* blood stages

Metabolic analysis of 3D7^Δ*PflipB*^ mutants highlighted an increased glycolytic activity, which is the main energy releasing pathway in blood stage *P. falciparum* (Salcedo-Sora et al., 2014). Increased glycolytic activity in 3D7^Δ*PflipB*^ mutants was mostly evident through increased glucose demand and increased lactate production via spent medium analysis (Fig. 3).

A potential model tying these phenotypes together goes as follows: changes to apicoplast redox balance due to depletion of LA also induce changes in cytosolic redox through plastid-to-cytosol signalling. This signal leads to temporary accelerated differentiation, which results in a higher demand on glycolysis. However, we cannot rule out that the changes in cytosolic redox regulator expression and asexual cycle progression are the result of the metabolic changes taking place in this mutant and not the cause. The uncertainty stems from the potential dual roles of LA in i/ regulating redox and ii/ supporting PDC dependent fatty acid synthesis. However, accumulating evidence suggests that the latter is not essential during intra-erythrocytic development (Cobbold et al., 2013) unless fatty acid starvation occurs (Botté et al., 2013). We thus propose that the changes in carbon metabolism observed upon *lipB* deletion are not the consequence of disruption of fatty acid metabolism. This hypothesis is further supported by the absence of similar carbon metabolic variations in the 3D7^Δ*Pfae3*^ mutant (Fig. S2), which lacks a fully-functional PDC due to the deletion of aE3, and which has only modest changes in cytosolic redox (Laine et al., 2015).

### How is *lipB* deletion influencing the TCA cycle?

The main change we detected in the TCA cycle using targeted metabolomics analysis was in the labelling of citrate, the first product of the cycle. In 3D7^Δ*PflipB*^ mutants, citrate presented mainly as M+2 labelling and showed a significant decrease in M+5 while all other isotopologues for this metabolite were below detection levels (Fig. 4C). These results point to a change in the cycle flow, whereby unlabelled glutamate and M+2 pyruvate are likely generating unlabelled and M+2 labelled citrate (Fig 4A). Furthermore, the apparent absence of M+4 and M+6 citrate fractions may suggest a non-cyclic pathway that starts with 2-oxoglutarate from glutamine and stops at citrate. The increase in relative intracellular levels for 2-oxoglutarate and succinate further support that the cycle is fuelled mainly by the glutamate branch.

The link between *lipB* deletion and the changes seen in the TCA cycle is unclear, and the lack of change in ICDH protein levels (Fig. 5A) is puzzling. One possible explanation is that the enzymatic activity of ICDH, or indeed citrate synthase and aconitase, may be inhibited in response to the observed cellular redox changes. Studies on plant mitochondria highlight the role of thiol redox switches in adjusting mitochondrial function in light of external stresses (reviewed in Nietzel et al., 2017), and citrate synthase, aconitase and ICDH are all substrate for thiol based redox regulation (Yoshida et al., 2013; Schmidtmann et al., 2014; Yoshida and Hisabori, 2014).

Interestingly, alongside the altered levels of citrate labelling, we observed an increase in relative levels of BCKDH E2 (Fig. 5A) and no change in the total relative intracellular levels of citrate (Fig. 4B). Previous studies suggested that citrate could take part in a malate shuttle, whereby cytosolic citrate is used to generate oxaloacetate and acetyl-CoA, which in turn take part in carbon fixation and protein acetylation (Cobbold et al., 2013; Storm et al., 2014). The apparent absence of M+4 and M+6 citrate in 3D7^Δ*PflipB*^ mutants is in line with this scenario, and could suggest that the citrate fraction that is not cycling in the TCA cycle but may instead be channelled towards this malate shuttle. If true, this would result in increased availability of cytosolic acetyl-CoA, which in turn may affect histone acetylation and thus gene expression patterns (Cobbold et al., 2016). This hypothetical model could explain the altered progression through the cell cycle, which typically depends on very tight regulation of gene expression (Bozdech et al., 2003).

### LA biosynthesis is essential to complete sporogony

We identified a severe defect in sporozoite production in the NF54^Δ*PflipB*^ mutant (Table 1, Fig. 6,7), despite the fact that normal numbers of oocysts were produced. This observation raises the possibility that LA synthesis in the apicoplast is essential for sporoblast development, through a yet undefined mechanism. This finding is surprising considering the evidence from the rodent parasite *P. berghei*, where deletion of *lipB* caused no defect to the production of salivary gland sporozoites but showed a moderate defect in liver-stage development (Falkard et al., 2013). This is not the first example for such a discrepancy, PDC E1α is dispensable for mosquito development in the rodent malaria parasite *P. yoelii* but was necessary for sporozoite maturation in *P. falciparum* (Cobbold et al., 2013). Taken together these findings point to different dependency of human and rodent malaria on PDC enzymes for development in the mosquito. A possible reason for this difference may be the increased number of sporozoites produced per oocyst, which is four-fold higher in the human malaria parasites *P. falciparum* (Rungsiwongse and Rosenberg, 1991) than in rodent malaria parasites (Lindner et al., 2013; Shimizu et al., 2010). High sporozoite numbers in rodent malaria oocysts may require enhanced metabolism, which would depend on both PDC activity and a functional redox regulation network.

## Material and Methods

### Parasite culture and assessment of growth

#### - *P. falciparum* culturing and synchronization

*P. falciparum* 3D7 parasites (isolated in the Netherlands) were cultured in RPMI 1640 (Invitrogen) supplemented with 11 mM D-Glucose (Sigma Aldrich), 0.5% w/v AlbuMAX II (Invitrogen), 200 mM hypoxanthine (Sigma Aldrich), and 20 mg/ml gentamycin (PAA) in human erythrocytes at 5% haematocrit (Trager and Jensen, 1976). Parasites were cultured maintaining a reduced oxygen atmosphere (1% O_2_, 3% CO_2_ and 96% N_2_) and a constant temperature of 37°C (referred to as standard procedures). Parasitemias were determined by microscopy analysis of Giemsa-stained thin smears and synchronisations were performed following the sorbitol procedure (Lambros and Vanderberg, 1979). Tighter synchronisation was obtained by a combination of sorbitol treatment with magnetic-activated cell sorting (MACS) using LD columns (Miltenyi Biotech). Briefly, sorbitol-synchronised parasites were maintained in culture until they reached the early ring stage. Cultures were synchronised twice with sorbitol 6 hours apart and then cultured until they reached the late schizont stage. Schizonts were purified over a MACS column once cultures had reached a schizont:ring ratio of 1:2. Schizonts were then placed in culture for 1 hour with gentle shaking and cultures synchronised again using the sorbitol method to obtain highly synchronous ring stages with 1-hour synchrony. Highly-synchronous parasites were used for all time point experiments and RNA extractions, with triplicate cultures for each condition.

#### - Spent medium metabolite quantification

Analyses of D-glucose and L-lactate concentrations from spent culture medium samples were performed using the D-Glucose-HK and L-Lactic acid kits from Megazyme following manufacturer's protocol. Briefly, parasite cultures for each condition were synchronised using a double sorbitol treatment (Lambros and Vanderberg, 1979) with a 6 hour window, and split in triplicate cultures at the same parasitemia. At each time point, an aliquot of the culture was collected from each condition and erythrocytes were pelleted by centrifuging at 1000g for 5 min. The resulting supernatant was then stored at −20°C. For the enzymatic assay, parasite spent medium samples were diluted by 1:6 with double distilled water.

### Evaluation of antioxidant gene expression

#### - RT-qPCR

Highly synchronous parasites (see above) were cultured in triplicate for each condition until they reached 26, 30 and 34 hpi. Pellets of infected RBCs at 6-8% parasitemias were then washed three times in PBS and kept at −80°C. Nucleic acids were extracted using the RNeasy kit (QIAGEN). Contaminating DNA was removed using the Turbo DNA-free kit (Thermo Fisher Scientific). RNA samples were then reverse transcribed using the RETRO-script kit (Thermo Fisher Scientific). qPCR was performed using the Power SYBR Green Master Mix (Thermo Fisher Scientific) adding 20 ng of cDNA for each reaction and 300 nM of each primer (see Table S1). All reactions were run in a 7500 Real-Time PCR System (Applied Biosystems). The calculation of relative gene expression was performed using the ΔΔ(Ct) method (Livak and Schmittgen, 2001).

#### - Protein extraction and quantitative fluorescent western blot

For protein extraction, saponin-lysed parasite pellets were resuspended in 2D lysis buffer (100 mM Hepes pH 7.4, 5 mM MgCl2,10 mM EDTA, 0.5% (v/v) Triton X-100, 5 μg/ml RNase A, 1x complete protease inhibitor cocktail (Roche) in ddH_2_O water). Samples were subjected to three rounds of freeze-thaw in dry ice and incubated at 4°C for 5 min in a sonicated water bath. Samples were then centrifuged at 13000 g for 20 min at 4°C. Supernatants containing protein fractions were quantified using the Protein Assay kit (Bio-Rad), with bovine serum albumin used to generate a reference quantification curve.

Western blot analysis was performed by separating 20 μg of protein sample by SDS-PAGE with NuPage Novex 4–12% and 15% (w/v) Bis-Tris gels (Invitrogen). Separated proteins were transferred to Protran nitrocellulose membranes (Schleicher & Schuell) using a Transblot semi-dry transfer system (BioRad). Membranes were blocked with 5% (w/v) dried skimmed milk in PBS for 1-18 hours and incubated for 1 hour with two or more primary antibodies. Primary antibodies used for relative quantification included *P. falciparum* rabbit anti-actin antibody (1:12,000, loading control), *P. falciparum* rabbit anti-1-CysPx (1-Cys peroxiredoxin) antibody 1:50,000; *P. falciparum* rabbit anti-2-CysPx (2-Cys peroxiredoxin) antibody at 1:70,000; *P. falciparum* rabbit anti-BCKDH E2 antibody at 1:5,000; *P. falciparum* rabbit anti-isocitrate dehydrogenase antibody at 1:10,000 and *P. falciparum* rabbit anti-PDC E2 lipoyl domain antibody at 1:250. Membranes were washed three times in PBS containing 0.2% (v/v) Tween 20 and 2.5% (w/v) dried skimmed milk. Blots were then probed with IR dye-conjugated antibody (1:10,000, IRDye800CW goat anti-rabbit antibody; LI-COR Biosciences) for 1 hour and washed again twice. Membranes were loaded in an Odyssey SA scanner (LI-COR Biosciences) and fluorescent signal intensities quantified with the Image Studio software (LI-COR Biosciences). All antibodies were custom made by Eurogentec.

### Metabolomics experiment and analysis

#### - ^13^C-U-D-glucose labelling experiment setup

Parasites were cultured until parasitemias attained 6-8%. Two sorbitol treatments were performed at approximately 8 and 14 hpi. After synchronisation, triplicate parasites cultures were set using a medium where D-glucose was replaced with ^13^C-U-D-glucose (99%, CK Gas Products Ltd) and the haematocrit was set to 1%. A control culture of uninfected RBCs was prepared with the same conditions. All cultures were incubated for 20 hours following standard procedures until late trophozoites. At this point, parasite culture metabolism was rapidly quenched at 4°C in a bath of dry ice and 70% ethanol (Vincent and Barrett, 2015). Erythrocyte pellets were then obtained and washed in ice-cold PBS by centrifugation at 800g for 5 min at 4°C. Infected RBCs in the pellets were enriched using MACS LD column purification (Miltenyi Biotech) and a QuadroMACS magnet (Miltenyi Biotech) with all steps performed at 4°C. Enriched samples were quantified using a Neubauer cell counting chamber and a Scepter 2.0 Handheld Automated Cell Counter (Millipore) to have 2.0×10^8^ infected RBC per sample. The same number of uninfected RBCs was collected as a control. All the samples were then added to a solution of HPLC-grade chloroform:methanol:water (1:3:1; v/v/v), at concentration of 2×10^8^ parasites per 0.5 mL solution, incubated in ice in a sonicating water bath for 2 min and extracted for 1 hour at 4°C and 1500 rpm on an orbital shaker. After extraction, samples were centrifuged at 13000g for 20 min at 4°C, and supernatants transferred into glass mass spectrometry vials (Thermo) and stored at −80°C until LC-MS analysis.

#### - Liquid chromatography-mass spectrometry analyses

Metabolomics analyses were performed by liquid chromatography-mass spectrometry using an Ultimate 3000 LC system (Dionex, UK) connected to a Q Exactive HF Hybrid Quadrupole-Orbitrap mass spectrometer, (Thermo Fisher Scientific). The system was controlled by the software Chromeleon (Dionex, UK) and Xcalibur (Thermo Scientific), acquiring both positive and negative ionisation mode. Chromatographic separation was performed with a ZIC-pHILIC chromatography column (150 mm64.6 mm65 mm; Sequant, Uemå, Sweden) using a two solvent system consisting of solvent A: 20 mM ammonium carbonate and solvent B: acetonitrile. The table shows chromatographic conditions:

| Solvent A (%) | Solvent B (%) | Time (min) |
| --- | --- | --- |
| 20 | 80 | 0 |
| 80 | 20 | 30 |
| 95 | 5 | 31 |
| 95 | 5 | 35 |
| 20 | 80 | 36 |
| 20 | 80 | 46 |

#### - Metabolomic data analysis

Vendor-specific raw data were initially centroided and converted into the open format mzXML for subsequent processing. PeakML files (Scheltema et al., 2011) were hen generated by extracting the chromatographic peaks contained in the mzXML files using the detection algorithm from XCMS (Tautenhahn et al., 2008). The data processing pipeline mzMatch.R (Jankevics et al., 2012) was used to sort and combine all PeakML files corresponding to replicates and to exclude all non-reproducible data. Further steps of noise-filtering, gap-filling, and metabolite identification were performed on PeakML files utilising data obtained from metabolic standards run in parallel. For each metabolite of interest, the proportions of each isotopologue and its relative abundance in the sample were determined. The PeakML.Isotope.TargetedIsotopes function of mzMatch-ISO (Chokkathukalam et al., 2013) was used to scan the PeakML files for labelled metabolite quality and quantity. All metabolites of interest in this study were reliably identified by comparison of the chromatographic retention times and the m/z values with an authentic metabolic standard processed in parallel. These should be then considered as “identified compounds” or level 1 according to the Metabolomic Standard Initiative (Sumner et al., 2007). All metabolomics data was corrected for natural carbon isotope abundance and reagent impurity using the software IsoCor (Millard et al., 2019).

### Double cross-over deletion of *Pf*LipB in NF54 strain by *Cre-loxP* system

*P. falciparum* strain NF54 was cultured in complete medium containing RPMI 1640 salts and 10% heat-inactivated human serum (Graves et al., 1984). The strategy for double cross over deletion of *pflipB* using *Cre-lox*P is depicted in **Fig. S5**. Briefly, we employed a double crossover recombination strategy to generate parasite lines lacking a functional *pflipB* locus (PlasmoDB ID: PF3D7_0823600). A 0.5 kb fragment of *pflipB,* PCR-amplified from NF54 gDNA using primers p1/p2 (Table S1), was cloned into PCC1-*cdup*-h*dhfr*-DXO between the *SacII* and *AflII* sites and served as the homology region for the first cross-over (O’Neill et al., 2011). The second homologous fragment (0.5 kb) was PCR amplified using primers p3/p4 and cloned between *EcoRI* and *AvrII* sites to give rise to 8.5 kb PCC1-*cdup-*h*dhfr-*Δ*lipB* plasmid. 50μg of this plasmid was used to transfect NF54 strain by electroporation and transformed parasites were selected with 1.5 nM WR99210. Drug pressure was maintained until 7 days and parasites were then cultured in drug-free medium up to day 27 when the parasites reappeared. WR99210-resistant parasites were then treated with 1.0 µM 5-Fluorocystosine (in DMSO) to remove single crossover-integrated plasmid and episomal forms, both of which contain the suicidal *cdup* marker (Maier et al., 2006). To unmark the parasites, pTET-BSD-*Cre* (kind gift from Allan Cowman) was mobilized into recombinant NF54 and cultured in 2.5 µM blasticidin S-hydrochloride for 7 days, after which single-cell cloning was performed (Goodyer and Taraschi, 1997).

### Generation of *Pf*Δ*lipB* gametocytes and production of salivary gland sporozoites

Gametocytes were induced from *Pf*Δ*lipB* and WT NF54 parasite lines by multiple rounds of sub-culturing for 14-18 days with nutrient deprivation, as described (Ponnudurai et al., 1986). Gametocyte induction and maturation was monitored microscopically by Giemsa-stained thin blood smears from culture samples. Mature gametocyte pellets were mixed with fresh O-type Rh^+^ blood and human serum to produce an artificial blood meal at approximately 50% hematocrit for mosquito feeds. The final concentration of stage V gametocytes in artificial blood meals was 1.0%±0.1% (mean±SEM across 7 experiments), which was equivalent to the gametocyte numbers observed with NF54 (**Table S1**). *Anopheles stephensi* mosquitoes were fed 3-5 days after their emergence from pupae. Fed mosquitoes were maintained for 13–16 days before recovery of salivary gland sporozoites (SPZ) by hand dissection.

### Ultra-structure analysis of midgut oocysts

The infected midgut oocysts were fixed in 4%/0.1% formaldehyde/glutaraldehyde and immersed in a 50:50 glycerol/water solution for Differential Interference Contrast (DIC) imaging. Using a Nikon Ti Eclipse inverted microscope, cut midgut sections were pre-screened for the presence of oocysts at 10× magnification using an automated tiling feature of the Nikon NIS Elements 3.2 software. Individual oocysts were imaged at high magnification using a 60× NA 1.4 oil immersion objective with a matching DIC slider. Images were compiled using the NIH's ImageJ program.

Morphology of developing oocysts (Fig 6A) was done by fixing dissected mosquito midguts in 2.5% glutaraldehyde in PBS, and then in 0.5%OsO4, dehydration in an ethanol series, embedding in London Resin White, and semi-thin sections (400nm) were mounted on glass slides and stained with 0.5% toluidine blue (w/v): 0.1% Na2CO3 (w/v) for 10 seconds, and then imaged on an Olympus BH-2 light microscope.

## Acknowledgements

We would like to thank Dr Sujaan Das, Dr Mahmood Alam, Dr Lewis King and Dr Sonal Sethia for their help and advice. The research was supported by the European Community’s Seventh Framework Programme [grant number FP7/2007-2013] under grant agreements No 242095 and No ParaMet 290080 (MB), by Medical Research Council grant number MR/S024573/1 (LS), and by the NIH (R01 AI085584 to DAF). The Wellcome Centre for Integrative Parasitology is supported by core funding from the Wellcome Trust [104111]. Finally, GIM was awarded Laureate Fellowship from the Australian Research Council and LS is a Royal Society of Edinburgh Personal Research Fellow.

## Supplemental figures

**Figure S1.**
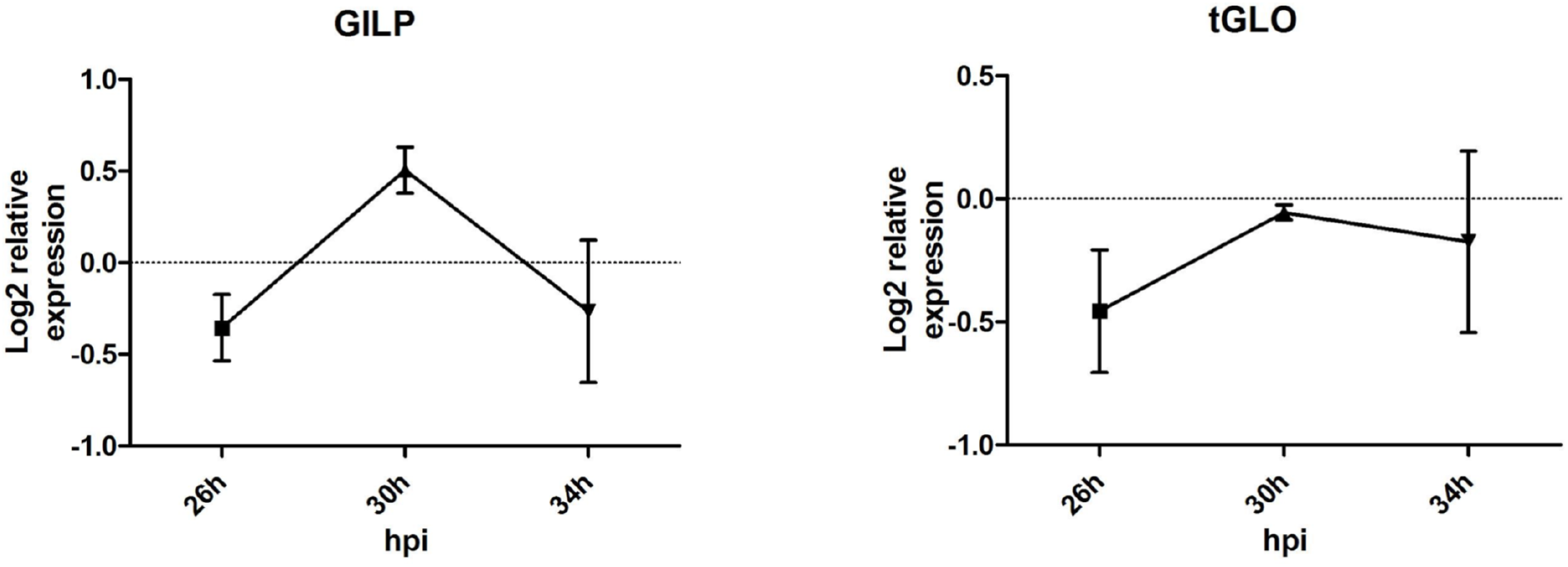
Analysis of the apicoplast glyoxalase system enzymes relative expression levels. Relative expression levels for the enzymes glyoxalase-1-like protein (GILP) and glyoxalase 2 (tGLO) were determined in three independent experiments from highly synchronised parasites following the sorbitol and MACS protocol (see Materials and Methods). Samples were harvested at 26, 30 and 34 hpi. Differences are expressed as Log_2_ of the 3D7^Δ*PflipB*^/3D7^WT^ ratio of the mean signals from three experiments ± SD.

**Figure S2.**
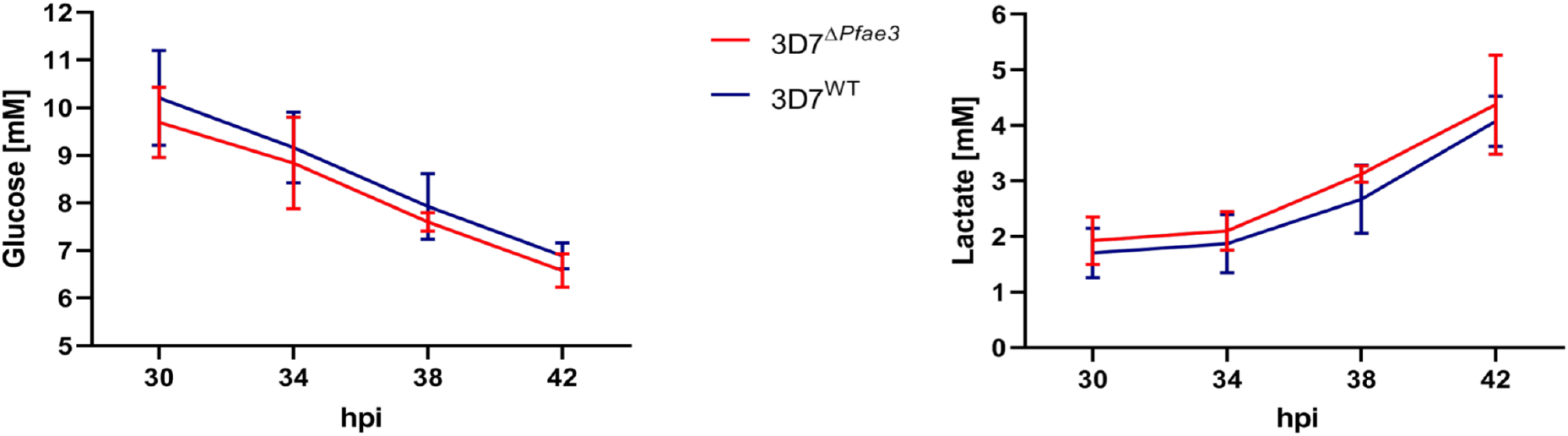
Analysis of D-glucose (left) and L-lactate (right) in spent medium samples from 3D7^Δ*Pfae3*^ mutants and 3D7^WT^ parasites cultures. Spent medium samples were analysed using a commercial enzymatic assay for D-glucose and L-lactate. Three independent experiments in triplicate cultures at 4% parasitemia were used for each parasite line tested in this experiment. Results are reported as mean ± SD (n=3). The variance between the lines at each time point was analysed with the Student t-test using GraphPad Prism 5.

**Figure S3.**
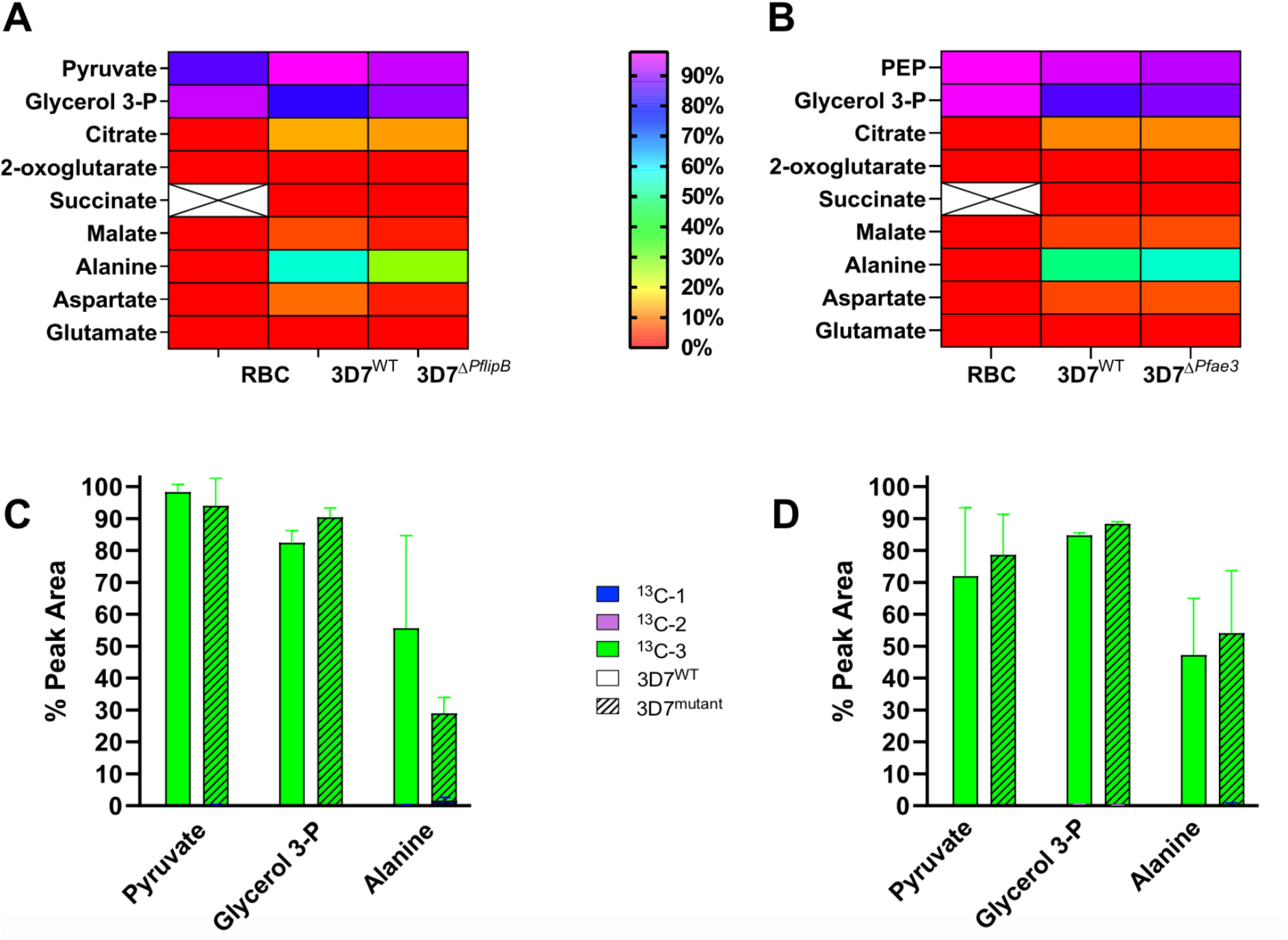
Metabolomic analyses of 3D7^Δ*PflipB*^ (A,C) 3D7^Δ*Pfae3*^ (B,D) mutants and 3D7^WT^ parasites using ^13^C-U-D-glucose labelling. Results from two independent targeted metabolomics experiments in biological triplicates comparing 3D7^Δ*Pfae3*^ or 3D7^Δ*PflipB*^ mutant to 3D7^WT^ line after incubation in culture medium containing 100% ^13^C-U-D-glucose for 28 hours. (**A-B**) Heatmap representing the total labelling incorporation from ^13^C-U-D-glucose in RBC, 3D7^WT^ and 3D7^Δ*PflipB*^ **(A)** or 3D7^Δ*Pfae3*^ **(B)** mutants. Crossed squares shows that succinate could not be detected in that analysis. (**C-D**) Bar graphs summarising the percentage of isotopic incorporation in each identified metabolite relative to the peak area presented as mean ± SD (n=2). Empty bars represent metabolites identified in 3D7^WT^ parasites, while dashed bars correspond to metabolites from 3D7^Δ*PflipB*^ **(C)** or 3D7^Δ*Pfae3*^ **(D)** mutants.

**Figure S4.**
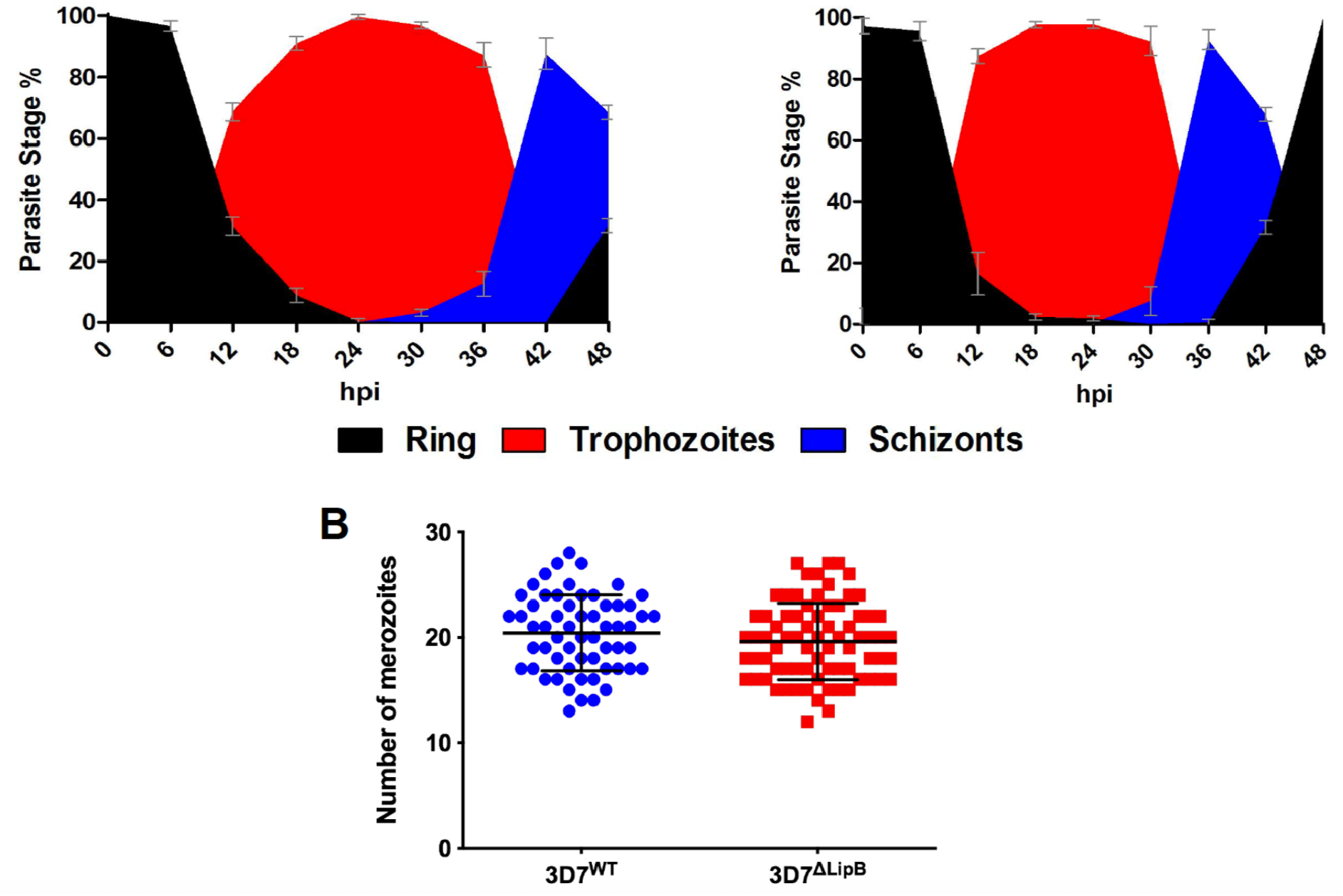
Growth pattern of 3D7^Δ*PflipB*^ mutants and 3D7^WT^ parasites. (**A**) The variation for each parasite stage during asexual development was estimated by counting 200 random infected RBCs for each time point for 3D7^WT^ (left) and 3D7^Δ*PflipB*^ mutants (right). Fractions are presented as the mean percentage from three cultures for each condition, error bars correspond to SD. (**B**) The graph represents the number of merozoite per segmenter (n=100) in 3D7^Δ*PflipB*^ mutants and 3D7^WT^. Bars represent mean ± SD.

**Figure S5.**
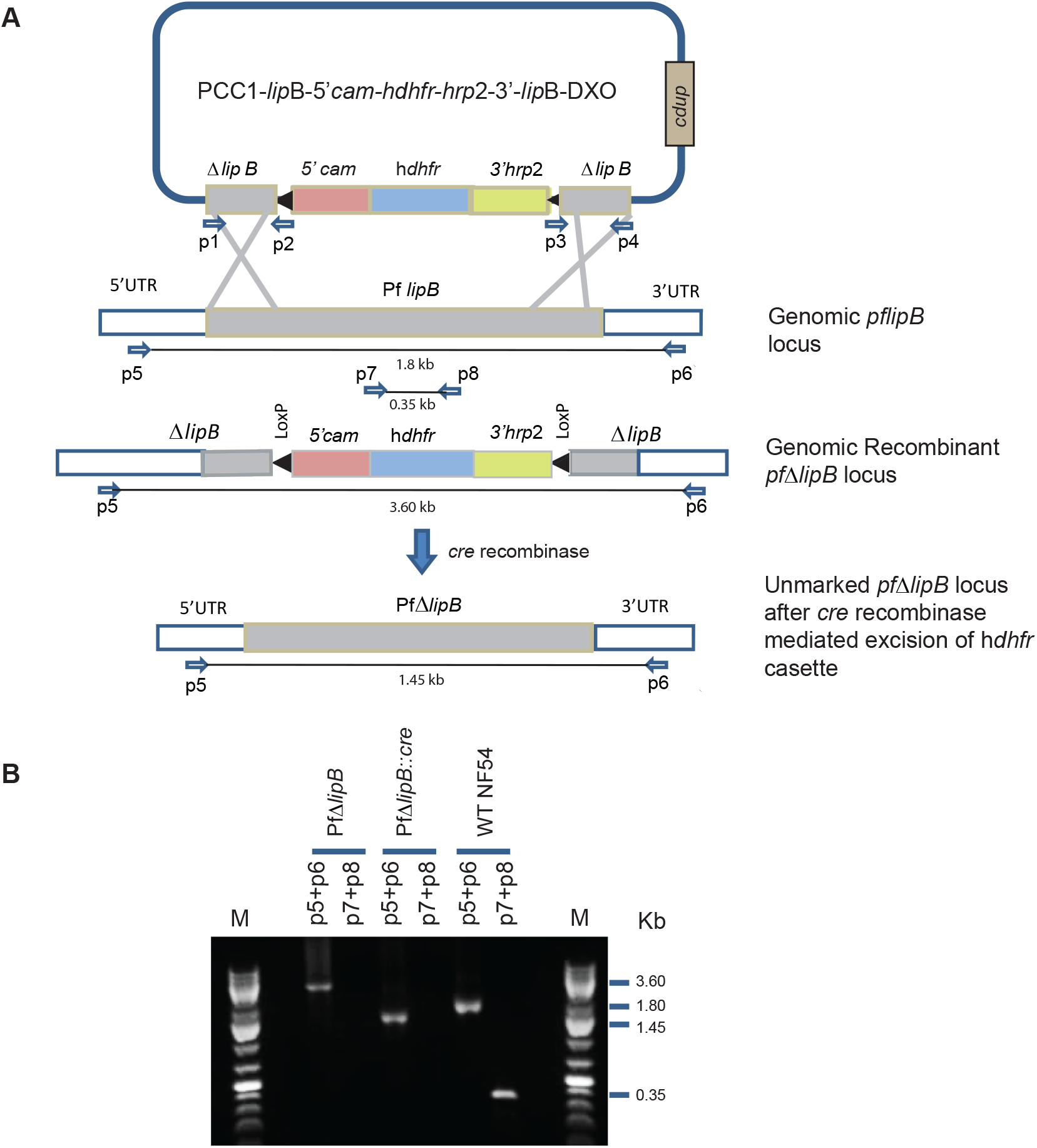
Generation and genotypic analysis of Pf *ΔlipB* in *P. falciparum* NF54. **(A)** To generate parasite lines lacking a functional *pflipB* locus (PF3D7_0823600) we employed a double crossover recombination strategy using *Cre-loxP*. 0.5 kb fragments on both ends of *pflipB* served as homology regions for double cross-over recombination and the LoxP site was incorporated into the genomic locus that flanks the h*dhfr* cassette. Primers used to amplify the double-recombination plasmid and to test for integration are shown. (**B**) PCR results showing *pflipB* deletion in the knockout strain. Lane 1: Marker (M) NEB 1kb plus DNA ladder, 3-4: *Pf*Δ*lipB*; Lanes 5-6: *Pf*Δ*lipB*::*Cre*, Lanes 7-8: wild-type NF54. Primers p5/p6 amplified a 3.6 kb fragment only from *Pf*Δ*lipB* strain due to the incorporation of the h*dhfr* cassette. A much smaller 1.45 kb fragment in *PfΔlipB::Cre* strain due to unmarking action of *Cre* recombinase. In both these strains primers p7/p8 did not give a product. Alternatively, primers p5/p6 amplified a 1.8 kb fragment in the wild-type NF54 strain. Primers p7/p8 amplified a product of 0.35 kb as the deleted fragment was retained in the non-recombinant WT NF54 strain.

**Table S1:**
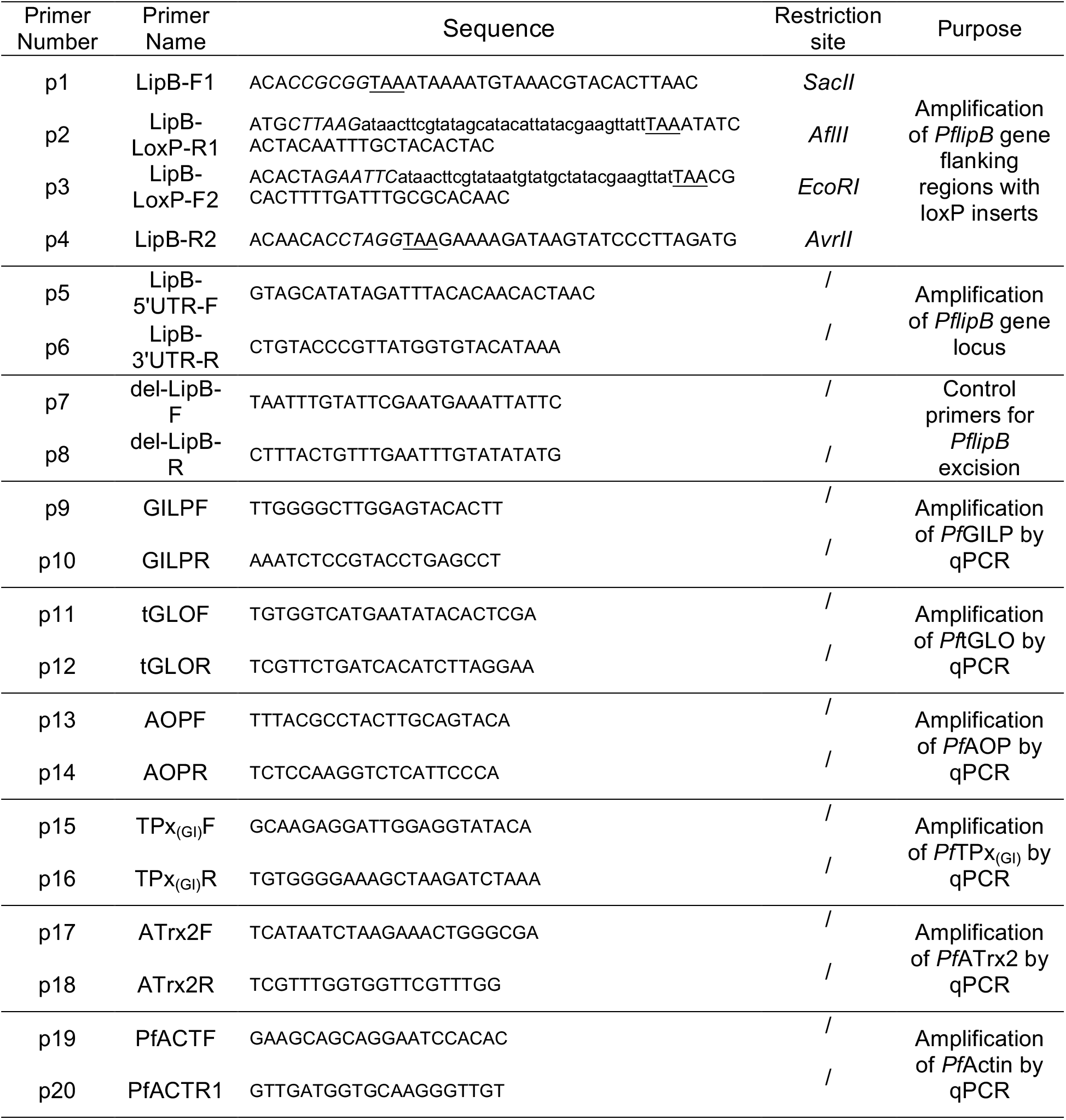
List of primers used in this work including names, sequences and purpose. Restriction enzyme sites are italicized. LoxP site are indicated in lower case letters. The underlined bases constitute STOP codon.

## Notes

### Competing Interest Statement

The authors have declared no competing interest.

## References

Akide-Ndunge, O.B., Tambini, E., Giribaldi, G., McMillan, P.J., Müller, S., Arese, P., Turrini, F., 2009. Co-ordinated stage-dependent enhancement of *Plasmodium falciparum* antioxidant enzymes and heat shock protein expression in parasites growing in oxidatively stressed or G6PD-deficient red blood cells. Malar. J. 8, 113. doi:10.1186/1475-2875-8-113

Becker, K., Tilley, L., Vennerstrom, J.L., Roberts, D., Rogerson, S., Ginsburg, H., 2004. Oxidative stress in malaria parasite-infected erythrocytes: Host-parasite interactions. Int. J. Parasitol. doi:10.1016/j.ijpara.2003.09.011

Biddau, M., Bouchut, A., Major, J., Saveria, T., Tottey, J., Oka, O., Van-Lith, M., Jennings, K.E., Ovciarikova, J., DeRocher, A., Striepen, B., Waller, R.F., Parsons, M., Sheiner, L., 2018. Two essential Thioredoxins mediate apicoplast biogenesis, protein import, and gene expression in *Toxoplasma gondii*. PLoS Pathog. 14, e1006836. doi:10.1371/journal.ppat.1006836

Biddau, M., Sheiner, L., 2019. Targeting the apicoplast in malaria. Biochem. Soc. Trans. 47, 973–983. doi:10.1042/BST20170563

Bilska, A., Włodek, L., 2005. Lipoic acid - the drug of the future? Pharmacol. Rep. 57, 570–7.

Bode, R., Ivanov, A.G., Hüner, N.P.A., 2016. Global transcriptome analyses provide evidence that chloroplast redox state contributes to intracellular as well as long-distance signalling in response to stress and acclimation in *Arabidopsis*. Photosynth. Res. 128, 287–312. doi:10.1007/s11120-016-0245-y

Botté, C.Y., Yamaryo-Botté, Y., Rupasinghe, T.W.T., Mullin, K.A., MacRae, J.I., Spurck, T.P., Kalanon, M., Shears, M.J., Coppel, R.L., Crellin, P.K., Maréchal, E., McConville, M.J., McFadden, G.I., 2013. Atypical lipid composition in the purified relict plastid (apicoplast) of malaria parasites. Proc. Natl. Acad. Sci. U. S. A. 110, 7506–11. doi:10.1073/pnas.1301251110

Boucher, M.J., Ghosh, S., Zhang, L., Lal, A., Jang, S.W., Ju, A., Zhang, S., Wang, X., Ralph, S.A., Zou, J., Elias, J.E., Yeh, E., 2018. Integrative proteomics and bioinformatic prediction enable a high-confidence apicoplast proteome in malaria parasites. PLoS Biol. 16, e2005895. doi:10.1371/journal.pbio.2005895

Bozdech, Z., Llinás, M., Pulliam, B.L., Wong, E.D., Zhu, J., DeRisi, J.L., 2003. The transcriptome of the intraerythrocytic developmental cycle of *Plasmodium falciparum*. PLoS Biol. 1, E5. doi:10.1371/journal.pbio.0000005

Bryk, R., Lima, C.D., Erdjument-Bromage, H., Tempst, P., Nathan, C., 2002. Metabolic enzymes of mycobacteria linked to antioxidant defense by a thioredoxin-like protein. Science 295, 1073–7. doi:10.1126/science.1067798

Bunik, V., Follmann, H., 1993. Thioredoxin reduction dependent on alpha-ketoacid oxidation by alpha-ketoacid dehydrogenase complexes. FEBS Lett. 336, 197–200. doi:10.1016/0014-5793(93)80801-Z

Bushell, E., Gomes, A.R., Sanderson, T., Anar, B., Girling, G., Herd, C., Metcalf, T., Modrzynska, K., Schwach, F., Martin, R.E., Mather, M.W., McFadden, G.I., Parts, L., Rutledge, G.G., Vaidya, A.B., Wengelnik, K., Rayner, J.C., Billker, O., 2017. Functional Profiling of a *Plasmodium* Genome Reveals an Abundance of Essential Genes. Cell 170, 260–272.e8. doi:10.1016/j.cell.2017.06.030

Chan, M., Sim, T.-S., 2005. Functional analysis, overexpression, and kinetic characterization of pyruvate kinase from Plasmodium falciparum. Biochem. Biophys. Res. Commun. 326, 188–96. doi:10.1016/j.bbrc.2004.11.018

Chokkathukalam, A., Jankevics, A., Creek, D.J., Achcar, F., Barrett, M.P., Breitling, R., 2013. mzMatch-ISO: an R tool for the annotation and relative quantification of isotope-labelled mass spectrometry data. Bioinformatics 29, 281–3. doi:10.1093/bioinformatics/bts674

Cobbold, S.A., Santos, J.M., Ochoa, A., Perlman, D.H., Llinás, M., 2016. Proteome-wide analysis reveals widespread lysine acetylation of major protein complexes in the malaria parasite. Sci. Rep. 6, 19722. doi:10.1038/srep19722

Cobbold, S.A., Vaughan, A.M., Lewis, I.A., Painter, H.J., Camargo, N., Perlman, D.H., Fishbaugher, M., Healer, J., Cowman, A.F., Kappe, S.H.I., Llinás, M., 2013. Kinetic flux profiling elucidates two independent acetyl-CoA biosynthetic pathways in *Plasmodium falciparum*. J. Biol. Chem. 288, 36338–50. doi:10.1074/jbc.M113.503557

Crawford, M.J., Thomsen-Zieger, N., Ray, M., Schachtner, J., Roos, D.S., Seeber, F., 2006. Toxoplasma gondii scavenges host-derived lipoic acid despite its de novo synthesis in the apicoplast. EMBO J. doi:10.1038/sj.emboj.7601189

Deponte, M., Becker, K., 2005. Glutathione S-transferase from malarial parasites: structural and functional aspects. Methods Enzymol. 401, 241–53. doi:10.1016/S0076-6879(05)01015-3

Dietz, K.-J., Turkan, I., Krieger-Liszkay, A., 2016. Redox- and Reactive Oxygen Species-Dependent Signaling into and out of the Photosynthesizing Chloroplast. Plant Physiol. 171, 1541–50. doi:10.1104/pp.16.00375

Dockrell, H.M., Playfair, J.H., 1984. Killing of *Plasmodium yoelii* by enzyme-induced products of the oxidative burst. Infect. Immun. 43, 451–6.

Falkard, B., Kumar, T.R.S., Hecht, L.-S., Matthews, K.A., Henrich, P.P., Gulati, S., Lewis, R.E., Manary, M.J., Winzeler, E.A., Sinnis, P., Prigge, S.T., Heussler, V., Deschermeier, C., Fidock, D.A., 2013. A key role for lipoic acid synthesis during Plasmodium liver stage development. Cell. Microbiol. 15, 1585–604. doi:10.1111/cmi.12137

Feeney, M.A., Veeravalli, K., Boyd, D., Gon, S., Faulkner, M.J., Georgiou, G., Beckwith, J., 2011. Repurposing lipoic acid changes electron flow in two important metabolic pathways of *Escherichia coli*. Proc. Natl. Acad. Sci. U. S. A. 108, 7991–6. doi:10.1073/pnas.1105429108

Foth, B.J., Stimmler, L.M., Handman, E., Crabb, B.S., Hodder, A.N., McFadden, G.I., 2005. The malaria parasite Plasmodium falciparum has only one pyruvate dehydrogenase complex, which is located in the apicoplast. Mol. Microbiol. 55, 39–53. doi:10.1111/j.1365-2958.2004.04407.x

Frohnecke, N., Klein, S., Seeber, F., 2015. Protein-protein interaction studies provide evidence for electron transfer from ferredoxin to lipoic acid synthase in *Toxoplasma gondii*. FEBS Lett. 589, 31–6. doi:10.1016/j.febslet.2014.11.020

Gamo, F.-J., Sanz, L.M., Vidal, J., de Cozar, C., Alvarez, E., Lavandera, J.-L., Vanderwall, D.E., Green, D.V.S., Kumar, V., Hasan, S., Brown, J.R., Peishoff, C.E., Cardon, L.R., Garcia-Bustos, J.F., 2010. Thousands of chemical starting points for antimalarial lead identification. Nature 465, 305–10. doi:10.1038/nature09107

Goodyer, I.D., Taraschi, T.F., 1997. *Plasmodium falciparum*: A Simple, Rapid Method for Detecting Parasite Clones in Microtiter Plates. Exp. Parasitol. 86, 158–160. doi:10.1006/EXPR.1997.4156

Gorąca, A., Huk-Kolega, H., Piechota, A., Kleniewska, P., Ciejka, E., Skibska, B., 2011. Lipoic acid - biological activity and therapeutic potential. Pharmacol. Rep. 63, 849–58. doi:10.1016/S1734-1140(11)70600-4

Graves, P.M., Carter, R., McNeill, K.M., 1984. Gametocyte production in cloned lines of *Plasmodium falciparum*. Am. J. Trop. Med. Hyg. 33, 1045–50. doi:10.4269/ajtmh.1984.33.1045

Guggisberg, A.M., Frasse, P.M., Jezewski, A.J., Kafai, N.M., Gandhi, A.Y., Erlinger, S.J., Odom John, A.R., 2018. Suppression of Drug Resistance Reveals a Genetic Mechanism of Metabolic Plasticity in Malaria Parasites. MBio 9, 1–16. doi:10.1128/mBio.01193-18

Günther, S., Storm, J., Müller, S., 2009. *Plasmodium falciparum*: organelle-specific acquisition of lipoic acid. Int. J. Biochem. Cell Biol. 41, 748–52. doi:10.1016/j.biocel.2008.10.028

Günther, S., Wallace, L., Patzewitz, E.-M., McMillan, P.J., Storm, J., Wrenger, C., Bissett, R., Smith, T.K., Müller, S., 2007. Apicoplast lipoic acid protein ligase B is not essential for *Plasmodium falciparum*. PLoS Pathog. 3, e189. doi:10.1371/journal.ppat.0030189

Haramaki, N., Han, D., Handelman, G.J., Tritschler, H.J., Packer, L., 1997. Cytosolic and mitochondrial systems for NADH- and NADPH-dependent reduction of alpha-lipoic acid. Free Radic. Biol. Med. 22, 535–42. doi:10.1016/S0891-5849(96)00400-5

Harris, M.T., Walker, D.M., Drew, M.E., Mitchell, W.G., Dao, K., Schroeder, C.E., Flaherty, D.P., Weiner, W.S., Golden, J.E., Morris, J.C., 2013. Interrogating a hexokinase-selected small-molecule library for inhibitors of *Plasmodium falciparum* hexokinase. Antimicrob. Agents Chemother. 57, 3731–7. doi:10.1128/AAC.00662-13

Harwaldt, P., Rahlfs, S., Becker, K., 2002. Glutathione S-transferase of the malarial parasite *Plasmodium falciparum*: characterization of a potential drug target. Biol. Chem. 383, 821–30. doi:10.1515/BC.2002.086

Heneberg, P., 2018. Redox Regulation of Hexokinases, Antioxidants & Redox Signaling. Mary Ann Liebert, Inc. 140 Huguenot Street, 3rd Floor New Rochelle, NY 10801 USA. doi:10.1089/ars.2017.7255

Jankevics, A., Merlo, M.E., de Vries, M., Vonk, R.J., Takano, E., Breitling, R., 2012. Separating the wheat from the chaff: a prioritisation pipeline for the analysis of metabolomics datasets. Metabolomics 8, 29–36. doi:10.1007/s11306-011-0341-0

Kagan, V.E., Shvedova, A., Serbinova, E., Khan, S., Swanson, C., Powell, R., Packer, L., 1992. Dihydrolipoic acid-a universal antioxidant both in the membrane and in the aqueous phase. Reduction of peroxyl, ascorbyl and chromanoxyl radicals. Biochem. Pharmacol. doi:10.1016/0006-2952(92)90482-X

Ke, H., Lewis, I.A., Morrisey, J.M., McLean, K.J., Ganesan, S.M., Painter, H.J., Mather, M.W., Jacobs-Lorena, M., Llinás, M., Vaidya, A.B., 2015. Genetic investigation of tricarboxylic acid metabolism during the *Plasmodium falciparum* life cycle. Cell Rep. 11, 164–74. doi:10.1016/j.celrep.2015.03.011

Kehr, S., Sturm, N., Rahlfs, S., Przyborski, J.M., Becker, K., 2010. Compartmentation of redox metabolism in malaria parasites. PLoS Pathog. 6, e1001242. doi:10.1371/journal.ppat.1001242

Kimata-Ariga, Y., Yuasa, S., Saitoh, T., Fukuyama, H., Hase, T., 2018. *Plasmodium*-specific basic amino acid residues important for the interaction with ferredoxin on the surface of ferredoxin-NADP+ reductase. J. Biochem. 164, 231–237. doi:10.1093/jb/mvy045

Krnajski, Z., Walter, R.D., Müller, S., 2001. Isolation and functional analysis of two thioredoxin peroxidases (peroxiredoxins) from *Plasmodium falciparum*. Mol. Biochem. Parasitol. 113, 303–308. doi:10.1016/S0166-6851(01)00219-5

Laine, L.M., 2014. Functional, biochemical and structural analyses of Plasmodium falciparum pyruvate dehydrogenase complex. PhD Thesis, University of Glasgow. University of Glasgow.

Laine, L.M., Biddau, M., Byron, O., Müller, S., 2015. Biochemical and structural characterization of the apicoplast dihydrolipoamide dehydrogenase of *Plasmodium falciparum*. Biosci. Rep. 35, 1–15. doi:10.1042/BSR20140150

Lambros, C., Vanderberg, J.P., 1979. Synchronization of *Plasmodium falciparum* erythrocytic stages in culture. J. Parasitol. 65, 418–20.

Liebau, E., Bergmann, B., Campbell, A.M., Teesdale-Spittle, P., Brophy, P.M., Lüersen, K., Walter, R.D., 2002. The glutathione S-transferase from *Plasmodium falciparum*. Mol. Biochem. Parasitol. 124, 85–90. doi:10.1016/S0166-6851(02)00160-3

Lim, L., McFadden, G.I., 2010. The evolution, metabolism and functions of the apicoplast. Philos. Trans. R. Soc. Lond. B. Biol. Sci. 365, 749–63. doi:10.1098/rstb.2009.0273

Lindner, S.E., Mikolajczak, S.A., Vaughan, A.M., Moon, W., Joyce, B.R., Sullivan, W.J., Kappe, S.H.I., 2013. Perturbations of *Plasmodium* Puf2 expression and RNA-seq of Puf2-deficient sporozoites reveal a critical role in maintaining RNA homeostasis and parasite transmissibility. Cell. Microbiol. 15, 1266–83. doi:10.1111/cmi.12116

Livak, K.J., Schmittgen, T.D., 2001. Analysis of relative gene expression data using real-time quantitative PCR and the 2(-Delta Delta C(T)) Method. Methods 25, 402–8. doi:10.1006/meth.2001.1262

MacRae, J.I., Dixon, M.W., Dearnley, M.K., Chua, H.H., Chambers, J.M., Kenny, S., Bottova, I., Tilley, L., McConville, M.J., 2013. Mitochondrial metabolism of sexual and asexual blood stages of the malaria parasite *Plasmodium falciparum*. BMC Biol. 11, 67. doi:10.1186/1741-7007-11-67

Maier, A.G., Braks, J.A.M., Waters, A.P., Cowman, A.F., 2006. Negative selection using yeast cytosine deaminase/uracil phosphoribosyl transferase in *Plasmodium falciparum* for targeted gene deletion by double crossover recombination. Mol. Biochem. Parasitol. 150, 118–21. doi:10.1016/j.molbiopara.2006.06.014

McMillan, P.J., Stimmler, L.M., Foth, B.J., McFadden, G.I., Müller, S., 2005. The human malaria parasite *Plasmodium falciparum* possesses two distinct dihydrolipoamide dehydrogenases. Mol. Microbiol. 55, 27–38. doi:10.1111/j.1365-2958.2004.04398.x

Millard, P., Delépine, B., Guionnet, M., Heuillet, M., Bellvert, F., Létisse, F., Wren, J., 2019. IsoCor: Isotope correction for high-resolution MS labeling experiments. Bioinformatics 35, 4484–4487. doi:10.1093/bioinformatics/btz209

Mohring, F., Pretzel, J., Jortzik, E., Becker, K., 2014. The redox systems of Plasmodium falciparum and *Plasmodium vivax*: comparison, in silico analyses and inhibitor studies. Curr. Med. Chem. 21, 1728–56.

Mohring, F., Rahbari, M., Zechmann, B., Rahlfs, S., Przyborski, J.M., Meyer, A.J., Becker, K., 2017. Determination of glutathione redox potential and pH value in subcellular compartments of malaria parasites. Free Radic. Biol. Med. 104, 104–117. doi:10.1016/j.freeradbiomed.2017.01.001

Mony, B.M., Mehta, M., Jarori, G.K., Sharma, S., 2009. Plant-like phosphofructokinase from Plasmodium falciparum belongs to a novel class of ATP-dependent enzymes. Int. J. Parasitol. 39, 1441–53. doi:10.1016/j.ijpara.2009.05.011

Mooney, B.P., Miernyk, J.A., Randall, D.D., 2002. The complex fate of alpha-ketoacids. Annu. Rev. Plant Biol. 53, 357–75. doi:10.1146/annurev.arplant.53.100301.135251

Moura, F.A., de Andrade, K.Q., dos Santos, J.C.F., Goulart, M.O.F., 2015. Lipoic Acid: its antioxidant and anti-inflammatory role and clinical applications. Curr. Top. Med. Chem. 15, 458–83. doi:10.2174/1568026615666150114161358

Müller, S., 2015. Role and Regulation of Glutathione Metabolism in *Plasmodium falciparum*. Molecules 20, 10511–34. doi:10.3390/molecules200610511

Nepveu, F., Turrini, F., 2013. Targeting the redox metabolism of Plasmodium falciparum. Future Med. Chem. doi:10.4155/fmc.13.159

Nietzel, T., Mostertz, J., Hochgräfe, F., Schwarzländer, M., 2017. Redox regulation of mitochondrial proteins and proteomes by cysteine thiol switches. Mitochondrion 33, 72–83. doi:10.1016/j.mito.2016.07.010

O’Neill, M.T., Phuong, T., Healer, J., Richard, D., Cowman, A.F., O’Neill, M.T., Phuong, T., Healer, J., Richard, D., Cowman, A.F., 2011. Gene deletion from *Plasmodium falciparum* using FLP and Cre recombinases: Implications for applied site-specific recombination. Int. J. Parasitol. 41, 117–123.

Oppenheim, R.D., Creek, D.J., Macrae, J.I., Modrzynska, K.K., Pino, P., Limenitakis, J., Polonais, V., Seeber, F., Barrett, M.P., Billker, O., McConville, M.J., Soldati-Favre, D., 2014. BCKDH: The Missing Link in Apicomplexan Mitochondrial Metabolism Is Required for Full Virulence of *Toxoplasma gondii* and *Plasmodium berghei*. PLoS Pathog. doi:10.1371/journal.ppat.1004263

Packer, L., Witt, E.H., Tritschler, H.J., 1995. alpha-Lipoic acid as a biological antioxidant. Free Radic. Biol. Med. 19, 227–50. doi:10.1016/0891-5849(95)00017-R

Pastrana-Mena, R., Dinglasan, R.R., Franke-Fayard, B., Vega-Rodríguez, J., Fuentes-Caraballo, M., Baerga-Ortiz, A., Coppens, I., Jacobs-Lorena, M., Janse, C.J., Serrano, A.E., 2010. Glutathione reductase-null malaria parasites have normal blood stage growth but arrest during development in the mosquito. J. Biol. Chem. 285, 27045–27056. doi:10.1074/jbc.M110.122275

Patzewitz, E.-M., Salcedo-Sora, J.E., Wong, E.H., Sethia, S., Stocks, P.A., Maughan, S.C., Murray, J.A.H., Krishna, S., Bray, P.G., Ward, S.A., Müller, S., 2013. Glutathione Transport: A New Role for PfCRT in Chloroquine Resistance. Antioxid. Redox Signal. doi:10.1089/ars.2012.4625

Perham, R.N., 2000. Swinging arms and swinging domains in multifunctional enzymes: catalytic machines for multistep reactions. Annu. Rev. Biochem. 69, 961–1004. doi:10.1146/annurev.biochem.69.1.961

Pick, U., Haramaki, N., Constantinescu, A., Handelman, G.J., Tritschler, H.J., Packer, L., 1995. Glutathione reductase and lipoamide dehydrogenase have opposite stereospecificities for alpha-lipoic acid enantiomers. Biochem. Biophys. Res. Commun. 206, 724–30. doi:10.1006/bbrc.1995.1102

Ponnudurai, T., Lensen, A.H.W., Meis, J.F.G.M., Meuwissen, J.H.E., 1986. Synchronization of *Plasmodium falciparum* gametocytes using an automated suspension culture system. Parasitology 93 (Pt 2), 263–74. doi:10.1017/S003118200005143X

Rahbari, M., Rahlfs, S., Przyborski, J.M., Schuh, A.K., Hunt, N.H., Fidock, D.A., Grau, G.E., Becker, K., 2017. Hydrogen peroxide dynamics in subcellular compartments of malaria parasites using genetically encoded redox probes. Sci. Rep. doi:10.1038/s41598-017-10093-8

Rungsiwongse, J., Rosenberg, R., 1991. The Number of Sporozoites Produced by Individual Malaria Oocysts. Am. J. Trop. Med. Hyg. 45, 574–577. doi:10.4269/ajtmh.1991.45.574

Salcedo-Sora, J.E., Caamano-Gutierrez, E., Ward, S.A., Biagini, G.A., 2014. The proliferating cell hypothesis: A metabolic framework for Plasmodium growth and development. Trends Parasitol. doi:10.1016/j.pt.2014.02.001

Scheltema, R.A., Jankevics, A., Jansen, R.C., Swertz, M.A., Breitling, R., 2011. PeakML/mzMatch: a file format, Java library, R library, and tool-chain for mass spectrometry data analysis. Anal. Chem. 83, 2786–93. doi:10.1021/ac2000994

Schmidtmann, E., König, A.-C., Orwat, A., Leister, D., Hartl, M., Finkemeier, I., 2014. Redox regulation of *Arabidopsis* mitochondrial citrate synthase. Mol. Plant 7, 156–69. doi:10.1093/mp/sst144

Seeber, F., Aliverti, A., Zanetti, G., 2005. The Plant-Type Ferredoxin-NADP+ Reductase/Ferredoxin Redox System as a Possible Drug Target Against Apicomplexan Human Parasites. Curr. Pharm. Des. 11, 3159–3172. doi:10.2174/1381612054864957

Seeber, F., Soldati-Favre, D., 2010. Metabolic pathways in the apicoplast of apicomplexa. Int. Rev. Cell Mol. Biol. 281, 161–228. doi:10.1016/S1937-6448(10)81005-6

Sewelam, N., Jaspert, N., Van Der Kelen, K., Tognetti, V.B., Schmitz, J., Frerigmann, H., Stahl, E., Zeier, J., Van Breusegem, F., Maurino, V.G., 2014. Spatial H2O2 signaling specificity: H2O2 from chloroplasts and peroxisomes modulates the plant transcriptome differentially. Mol. Plant 7, 1191–210. doi:10.1093/mp/ssu070

Sheiner, L., Demerly, J.L., Poulsen, N., Beatty, W.L., Lucas, O., Behnke, M.S., White, M.W., Striepen, B., 2011. A systematic screen to discover and analyze apicoplast proteins identifies a conserved and essential protein import factor. PLoS Pathog. 7, e1002392. doi:10.1371/journal.ppat.1002392

Sheiner, L., Vaidya, A.B., McFadden, G.I., 2013. The metabolic roles of the endosymbiotic organelles of *Toxoplasma* and *Plasmodium* spp. Curr. Opin. Microbiol. 16, 452–8. doi:10.1016/j.mib.2013.07.003

Shimizu, S., Osada, Y., Kanazawa, T., Tanaka, Y., Arai, M., 2010. Suppressive effect of azithromycin on Plasmodium berghei mosquito stage development and apicoplast replication. doi:10.1186/1475-2875-9-73

Shivapurkar, R., Hingamire, T., Kulkarni, A.S., Rajamohanan, P.R., Reddy, D.S., Shanmugam, D., 2018. Evaluating antimalarial efficacy by tracking glycolysis in Plasmodium falciparum using NMR spectroscopy. Sci. Rep. 8, 1–10. doi:10.1038/s41598-018-36197-3

Siciliano, G., Santha Kumar, T.R., Bona, R., Camarda, G., Calabretta, M.M., Cevenini, L., Davioud-Charvet, E., Becker, K., Cara, A., Fidock, D.A., Alano, P., 2017. A high susceptibility to redox imbalance of the transmissible stages of *Plasmodium falciparum* revealed with a luciferase-based mature gametocyte assay. Mol. Microbiol. 104, 306–318. doi:10.1111/mmi.13626

Storm, J., Sethia, S., Blackburn, G.J., Chokkathukalam, A., Watson, D.G., Breitling, R., Coombs, G.H., Müller, S., 2014. Phosphoenolpyruvate carboxylase identified as a key enzyme in erythrocytic *Plasmodium falciparum* carbon metabolism. PLoS Pathog. 10, e1003876. doi:10.1371/journal.ppat.1003876

Sumner, L.W., Amberg, A., Barrett, D., Beale, M.H., Beger, R., Daykin, C.A., Fan, T.W.M., Fiehn, O., Goodacre, R., Griffin, J.L., Hankemeier, T., Hardy, N., Harnly, J., Higashi, R., Kopka, J., Lane, A.N., Lindon, J.C., Marriott, P., Nicholls, A.W., Reily, M.D., Thaden, J.J., Viant, M.R., 2007. Proposed minimum reporting standards for chemical analysis: Chemical Analysis Working Group (CAWG) Metabolomics Standards Initiative (MSI). Metabolomics. doi:10.1007/s11306-007-0082-2

Tautenhahn, R., Bottcher, C., Neumann, S., 2008. Highly sensitive feature detection for high resolution LC/MS. BMC Bioinformatics. doi:10.1186/1471-2105-9-504

Tibullo, D., Li Volti, G., Giallongo, C., Grasso, S., Tomassoni, D., Anfuso, C.D., Lupo, G., Amenta, F., Avola, R., Bramanti, V., 2017. Biochemical and clinical relevance of alpha lipoic acid: antioxidant and anti-inflammatory activity, molecular pathways and therapeutic potential. Inflamm. Res. 66, 947–959. doi:10.1007/s00011-017-1079-6

Trager, W., Jensen, J.B., 1976. Human malaria parasites in continuous culture. Science 193, 673–5. doi:10.1038/098448b0

Urscher, M., Przyborski, J.M., Imoto, M., Deponte, M., 2010. Distinct subcellular localization in the cytosol and apicoplast, unexpected dimerization and inhibition of *Plasmodium falciparum* glyoxalases. Mol. Microbiol. 76, 92–103. doi:10.1111/j.1365-2958.2010.07082.x

van Schaijk, B.C.L., Kumar, T.R.S., Vos, M.W., Richman, A., van Gemert, G.-J., Li, T., Eappen, A.G., Williamson, K.C., Morahan, B.J., Fishbaugher, M., Kennedy, M., Camargo, N., Khan, S.M., Janse, C.J., Sim, K.L., Hoffman, S.L., Kappe, S.H.I., Sauerwein, R.W., Fidock, D.A., Vaughan, A.M., 2014. Type II fatty acid biosynthesis is essential for *Plasmodium falciparum* sporozoite development in the midgut of *Anopheles* mosquitoes. Eukaryot. Cell 13, 550–9. doi:10.1128/EC.00264-13

Vaughan, A.M., O’Neill, M.T., Tarun, A.S., Camargo, N., Phuong, T.M., Aly, A.S.I., Cowman, A.F., Kappe, S.H.I., 2009. Type II fatty acid synthesis is essential only for malaria parasite late liver stage development. Cell. Microbiol. 11, 506–20. doi:10.1111/j.1462-5822.2008.01270.x

Vincent, I.M., Barrett, M.P., 2015. Metabolomic-based strategies for anti-parasite drug discovery. J. Biomol. Screen. doi:10.1177/1087057114551519

Wezena, C.A., Alisch, R., Golzmann, A., Liedgens, L., Staudacher, V., Pradel, G., Deponte, M., 2017. The cytosolic glyoxalases of *Plasmodium falciparum* are dispensable during asexual blood-stage development. Microb. cell (Graz, Austria) 5, 32–41. doi:10.15698/mic2018.01.608

World Health Organization, 2019. World Malaria Report 2019. Geneva.

Wrenger, C., Müller, S., 2003. Isocitrate dehydrogenase of *Plasmodium falciparum*: Energy metabolism or redox control? Eur. J. Biochem. doi:10.1046/j.1432-1033.2003.03536.x

Yoshida, K., Hisabori, T., 2014. Mitochondrial isocitrate dehydrogenase is inactivated upon oxidation and reactivated by thioredoxin-dependent reduction in Arabidopsis. Front. Environ. Sci. 2, 1–7. doi:10.3389/fenvs.2014.00038

Yoshida, K., Noguchi, K., Motohashi, K., Hisabori, T., 2013. Systematic exploration of thioredoxin target proteins in plant mitochondria. Plant Cell Physiol. doi:10.1093/pcp/pct037

Zhang, M., Wang, C., Otto, T.D., Oberstaller, J., Liao, X., Adapa, S.R., Udenze, K., Bronner, I.F., Casandra, D., Mayho, M., Brown, J., Li, S., Swanson, J., Rayner, J.C., Jiang, R.H.Y., Adams, J.H., 2018. Uncovering the essential genes of the human malaria parasite Plasmodium falciparum by saturation mutagenesis. Science 360, eaap7847. doi:10.1126/science.aap7847

Zocher, K., Fritz-Wolf, K., Kehr, S., Fischer, M., Rahlfs, S., Becker, K., 2012. Biochemical and structural characterization of *Plasmodium falciparum* glutamate dehydrogenase 2. Mol. Biochem. Parasitol. 183, 52–62. doi:10.1016/j.molbiopara.2012.01.007

Zuzarte-Luís, V., Mello-Vieira, J., Marreiros, I.M., Liehl, P., Chora, Â.F., Carret, C.K., Carvalho, T., Mota, M.M., 2017. Dietary alterations modulate susceptibility to *Plasmodium* infection. Nat. Microbiol. 2, 1600–1607. doi:10.1038/s41564-017-0025-2

